# Microglial activation in an amyotrophic lateral sclerosis-like model caused by *Ranbp2* loss and nucleocytoplasmic transport impairment in retinal ganglion neurons

**DOI:** 10.1101/504225

**Authors:** Kyoung-in Cho, Dosuk Yoon, Minzhong Yu, Neal S. Peachey, Paulo A. Ferreira

**Affiliations:** Department of Ophthalmology Duke University Medical Center, Durham, NC 27710; Department of Ophthalmic Research, Cole Eye Institute, Cleveland Clinic Foundation, Cleveland, OH 44195; Research Service, Louis Stokes Cleveland Veterans Affairs Medical Center, Cleveland, OH 44106; Department of Ophthalmology, Cleveland Clinic Lerner College of Medicine of Case Western Reserve University, Cleveland, OH 44195

**Author notes:** To whom correspondence should be addressed: Paulo A. Ferreira, Duke University Medical Center, DUEC 3802, 2351 Erwin Road, Durham, NC 27710.

**Keywords:** Ran-binding protein 2 (Ranbp2), amyotrophic lateral sclerosis (ALS), microglia, nucleocytoplasmic transport, metalloproteinase-28, acetyl-CoA carboxylase 1, retinal ganglion neurons

## Abstract

Nucleocytoplasmic transport is dysregulated in sporadic and familial amyotrophic lateral sclerosis (ALS) and retinal ganglion neurons (RGNs) are purportedly involved in ALS. The Ran-binding protein 2 (Ranbp2) controls rate-limiting steps of nucleocytoplasmic transport. Mice with *Ranbp2* loss in Thy1^+^-motoneurons develop cardinal ALS-like traits, but the impairments in RGNs and the degree of dysfunctional consonance between RGNs and motoneurons caused by *Ranbp2* loss are unknown. This understanding will facilitate to discern the role of nucleocytoplasmic transport in the differential vulnerability of neurons to ALS and to develop therapeutic approaches and biomarkers in ALS. Here, we ascertain Ranbp2’s function and endophenotypes in RGNs of an ALS-like mouse model lacking *Ranbp2* in motoneurons and RGNs. Thy1^+^-RGNs lacking *Ranbp2* shared with motoneurons the dysregulation of nucleocytoplasmic transport. RGN abnormalities were comprised morphologically by soma hypertrophy and optic nerve axonopathy and physiologically by a delay of the visual pathway’s evoked potentials. Whole-transcriptome analysis showed restricted transcriptional changes in optic nerves that were distinct from those found in sciatic nerves. Specifically, the level and nucleocytoplasmic partition of the anti-apoptotic and novel substrate of Ranbp2, Pttg1/securin, were dysregulated. Further, acetyl-CoA carboxylase 1, which modulates *de novo* synthesis of fatty acids and T-cell immunity, showed the highest up-regulation (35-fold). This effect was reflected by the activation of ramified Cd11b^+^ and CD45^+^-microglia, increase of F4\80^+^-microglia and a shift from pseudopodial/lamellipodial to amoeboidal F4\80^+^-microglia intermingled between RGNs of naive mice. This immunogenic phenotype was accompanied by the intracellular sequestration in RGNs of metalloproteinase-28, which regulates macrophage recruitment and polarization in inflammation. *Ranbp2* genetic insults in RGNs and motoneurons trigger distinct paracrine signaling likely by the dysregulation of nucleocytoplasmic transport of neural-type selective substrates. Metabolic and immune-modulators underpinning RGN-to-microglial signaling are regulated by Ranbp2, and this neuroglial system manifests endophenotypes that are likely useful in the prognosis and diagnosis of ALS.

## Introduction

A growing body of evidence supports that heterogeneous forms of sporadic and familial ALS share impairments of nucleocytoplasmic transport [1–9]. Emerging evidence indicate that these impairments drive also the pathogenesis of other neurodegenerative diseases [10–13]. Further, substrates of the Ran GTPase cycle, which is the central driver of nucleocytoplasmic transport, act as potent genetic modifiers of ALS. This is thought to result from changes in nuclear-cytoplasmic distribution of nuclear shuttling substrates that promote their proteotoxicity in ALS [7,8,14–16]. Ran GTPase is a master regulator of nucleocytoplasmic transport by imparting a Ran-GTP to Ran-GDP gradient between the nucleus and cytoplasm. Ran-GTP associates to nuclear export receptors (e.g. exportin-1/CRM1) with their cargoes and to nuclear import receptors (e.g. importin-β) in the nuclear compartment, whereas the destabilization and hydrolysis of Ran-GTP from these nuclear export ensembles promotes their disassembly after exiting the nuclear pore complex (NPCs) [17]. The Ran-binding protein 2 (Ranbp2; also called Nup358) is a unique vertebrate and peripheral nucleoporin, which comprises the cytoplasmic filaments emanating from NPCs [18,19]. Ranbp2 plays a central role in terminal steps of nuclear export. This is achieved by multi-step events, such as the destabilization of the binding of Ran-GTP from nuclear export ensembles (e.g. importin-β) via the tandem Ran-GTP-binding domains (RBDs) of Ranbp2 [18,20,21], the recruitment of SUMOylated-modified Ran GTPase-activating protein (RanGAP) to Ranbp2 for the stimulation of Ran-GTP hydrolysis [22–25] and the docking of exportin-1 ensembles to the zinc-finger domain of Ranbp2 [26–28]. Impairment of Ranbp2 and some of its modular activities promote the dysregulation of nucleocytoplasmic transport of nuclear import and export receptors and the mislocalization of accessory substrates of Ranbp2 in a cell type-dependent manner [27,29–34].

We have shown that conditional loss of *Ranbp2* in Thy1^+^-motoneurons of *SLICK-H::Ranbp2^flox/flox^* mice drives ALS-like motor traits that are manifested by rapid declines of motor performance and conduction velocity of sciatic nerve followed by hind limb paralysis and respiratory distress, which culminates with the death of mice [34]. These motor behaviors are accompanied by the delocalization in motoneurons of Ran GTPase, exportin-1, importin-β and nuclear shuttling of substrates, such as histone deacetylase 4 (HDAC4) [34]. In addition, differential transcriptome and expression analysis of sciatic nerves and cell bodies of spinal motoneurons between mice with and without *Ranbp2* in motoneurons revealed that loss of *Ranbp2* triggers the dysregulation of neuroglia and chemokine signaling without apparent gliosis [34].

Multiple lines of evidence indicate that ALS and Ranbp2 affect retinal ganglion neurons (RGNs). The nuclear envelopes of RGNs, like motoneurons, are highly enriched in nuclear pores and Ranbp2, a phenotype that appears related to the long-distance burden of transport of cargoes exiting the nucleus towards axons and synapses [35]. In the inner retina, the prolyl isomerase activity of the cyclophilin domain of Ranbp2 also controls the levels of hnRNPA2B1 [33], a substrate in which mutations cause ALS [36]. Further, motoneurons and RGNs co-opt disease-causing substrates, such as optineurin, in which different mutations lead either to ALS [37,38] or the heterogeneous and leading blindness disorder, glaucoma, which is caused by the dysfunction and ultimately degeneration of RGNs [39,40]. Parenthetically, a glaucoma-causing mutation in optineurin disrupts its nucleocytoplasmic trafficking [41]. Sporadic and familial forms of ALS appear to produce non-motor syndrome phenotypes in humans, such as functional changes in the evoked potential of the visual pathway as well as subtle structural changes in the inner retina that includes the thinning of the nerve fiber layer [6,42–45]. Notably, some of these changes may occur even before clinical motor symptoms ensue. Regardless, the pathogenic drivers shared by motoneurons and RGNs in neurological diseases, such as ALS, and the reasons of the selective vulnerability of these neurons to dysfunction and/or degeneration by mutations in ubiquitously expressed genes, are elusive [46]. This barrier also highlights our poor understanding of the pathobiology of diseases affecting motoneurons, RGNs or both, and hinders the potential of using RGNs as diagnostic and prognostic tools of ALS and other neurological diseases.

In light of the foregoing findings, herein we ascertain neural-type selective roles of Ranbp2 in Thy1^+^-RGNs (aka retinal ganglion cells) in a mouse model of ALS, *SLICK-H::Ranbp2^flox/flox^*, in which both motoneurons and RGNs lose *Ranbp2.* Loss of *Ranbp2* in Thy1^+^-RGNs also impairs nucleocytoplasmic transport in these neurons. This impairment was accompanied by a significant reduction in the caliber of myelinated axons of the optic nerve, but without loss of retinal ganglion cell bodies. Notably, loss of *Ranbp2* in RGNs elicited transcriptome changes in the optic nerve that are distinct from those uncovered previously in the sciatic nerve [34]. Transcriptome analysis of optic nerves led to the identification of novel substrates of Ranbp2, such as Pttg1/securin, and metabolic targets, such as acetyl-CoA carboxylase 1 *(Acc1)*, which is implicated by several studies at the crux of metabolic shifts and

T-cell immunomodulation. In this respect, we found that loss of *Ranbp2* in RGNs triggered the activation of microglia and the intracellular sequestration in somata of RGNs of metalloproteinase 28 (Mmp28), which is also another factor implicated in the control of inflammatory responses. Taken together, the results indicate that Ranbp2 modulates neuroimmunity by regulation of *Acc1* expression, and biogenesis and proteostasis of Mmp28. These effects culminated in the non-cell autonomous activation of microglia by loss of Ranbp2 in RGNs.

## Materials and methods

### Mice

The generation of *SLICK-H::Ranbp2^flox/flox^* with the conditional deletion of *Ranbp2* in motoneurons and RGNs has been described [34]. Briefly, *Thy1-cre/ER^T2^-EYFP* (*SLICK-H*) mice with expression of cre and YFP in Thy1^+^-motoneurons and RGNs [47] were crossed with *Ranbp2 ^flox/flox^* [32,48,49] to produce *SLICK-H::Ranbp2 ^flox/flox^.* The generation of transgenic mice expressing wild-type *Ranbp2* tagged with HA in a constitutive *null Ranbp2* background, *Tg-Ranbp2^HA^::Ranbp2^−/−^*, and of *Tg-Ranbp2^RBD2/3*-HA^* mice with mutations in RBD2 and RBD3 of Ranbp2, were previously described [32,33]. All transgenic lines were on a mixed genetic background and did not carry the *rd1* and *rd8* alleles. Tamoxifen (T5648; Sigma-Aldrich) was administered by oral gavage for 5 consecutive days (0.25 mg/g of body weight) to 4-6 week-old mice as described previously [34]. Mice were reared at <70 lux in a pathogen-free transgenic barrier facility at Duke University with a 12 h light-dark cycles (6:00 A.M. – 6:00 P.M.) under humidity- and temperature-controlled conditions. Mice were given *ad libitum* access to water and chow diet 5LJ5 (Purina, Saint Louis, MO). Mice of either sex were examined by this study. The Institutional Animal Care and Use Committee of Duke University (A003-14-01) and Cleveland Clinic (2011-0580) approved the mouse protocols and the experiments were performed in accordance with NIH guidelines for the care and use of laboratory animals.

### Fluorescein Fundoscopy

Eyes of mice were dilated with a drop of atropine sulfate ophthalmic solution (1%; Alcon Laboratories, Inc., Fort Worth, TX) and then after 5min with phenylephrine hydrochloride (10%; HUB Pharmaceuticals, LLC, Rancho Cucamonga, CA). To capture images of YFP-labeled RGNs *in vivo* in mice, non-invasive fundus images were taken with Micron III imaging system (Phoenix Research Laboratories, Pleasanton, CA) in which a Xenon light source is coupled to a CCD-camera coupled microscope with an image resolution of 4 μm in a field of view of 1.8 mm. The pictures were captured in recording mode for 60 s and then the best-focused frames were selected and extracted.

### Visual evoked potential

After overnight dark adaptation mice were anesthetized with ketamine (80 mg/kg) and xylazine (16 mg/kg), and placed on a temperature-regulated heating pad. The pupils were dilated with eyedrops (2.5% phenylephrine HCl, 1% cyclopentolate, 1% tropicamide). VEPs were recorded using an active electrode positioned subcutaneously along the midline of the visual cortex and referenced to a needle electrode placed over the temporal cortex while a second needle electrode was inserted in the tail to serve as the ground lead [33,50]. VEPs were recorded to achromatic strobe flash stimuli presented in the LKC (Gaithersburg, MD) ganzfeld under dark-adapted conditions. The interstimulus interval ranged from 1.1 to 6 s, increasing with stimulus luminance from −2.4 to 2.1 log cd s/m^2^. The amplifier band-pass was set at 1-100 Hz and up to 60 successive responses were averaged to obtain a single VEP waveform. The mouse VEP is dominated by a negative component, N1. The implicit time of the N1 component was measured at the negative peak. The amplitude of the VEP was measured to N1 from the preceding baseline or positive peak (P1).

### Transmission electron microscopy

Mice were anesthetized with ketamine/xylazine (100mg/kg and 10mg/kg of body weight, respectively) and then cardiacally perfused with 2.5% glutaraldehyde and 4% paraformaldehyde in 0.1 M sodium cacodylate buffer, pH 7.4. Eyeballs and optic nerves were fixed for 2 h at room temperature followed by 18 h at 4°C in the same fixative, post-fixed in OsO_4_, and embedded in Araldite. Ultrathin sections were stained with uranyl acetate and lead citrate. Specimens were examined on a Phillips BioTwin CM120 electron microscope equipped with Gatan Orius and Olympus Morada digital cameras.

### Antibodies

The following and previously characterized antibodies were used for immunofluorescence (IF) or immunoblots (IB): rabbit anti-Ranbp2 (8 μg/ml (IF), Ab-W1W2#10) [33,34], rabbit anti-hsc70 (1:3,000 (IB), ENZO Life Science, Farmingdale, NY, cat# ADI-SPA-816) [34]; mouse mAb414 against nuclear pore complex proteins Nup62, Nup153, and Ranbp2/Nup358 (10 μg/ml (IF), Covance, Emeryville, CA, cat# MMS-120P) [34,49]; rabbit anti-Ran-GTP (1:100 (IF), gift from Dr. Ian Macara) [34,51]; rabbit anti-CRM1 (1:50, (IF), Santa Cruz Biotechnology, Santa Cruz, CA, cat# sc-5595) [34]; mouse Ran GTPase (1:100 (IF), BD Biosciences, San Jose, CA, cat# 610341) [34]; mouse anti-importin β (Mab3E9, 1:100, (IF), gift from Dr. Steve Adams) [34,52]; rabbit anti-HDAC4 (1:500 (IF), Santa Cruz Biotechnology, cat# sc11418) [34]; rabbit anti-Mmp28 (1:100, (IF), 1:1,000 (IB), Proteintech, Rosemont, IL; Cat: 18237-1-AP) [34]; rabbit anti-GFAP (1:200, (IF), DAKO, Carpinteria, CA, cat# z0334) [34,53]; rat anti-CD11b (1:100, (IF), AbD Serotec, Raleigh, NC, cat# MCA275G) [53,34]; rat anti-CD45 (1:100, (IF), BD Biosciences (BD Pharmingen), cat# 550239) [53]; rat anti-F4/80 (1:100, (IF), AbD Serotec, cat# MCA9976A); mouse anti-α synuclein (1:100, (IF), Millipore, cat# 36-008-C); goat anti-Brn3 (1:50 (IF), Santa Cruz Biotechnology, cat# sc6026); rabbit anti-Pttg1 (1:100, (IF), 1:1000 (IB), Proteintech, cat# 18040-1-AP); rabbit anti-NR2F2 (1:100 (IF), Abcam) [54]; Alexa Fluor-conjugated secondary antibodies and Hoechst 33 342 were from Invitrogen (Carlsbad, CA).

### Immunohistochemistry

Mice were anesthetized with ketamine/xylazine (100 mg/kg and 10 mg/kg of body weight, respectively) followed by cardiac perfusion with 2% paraformaldehyde in 1xPBS. Eye balls were dissected and incubated with 2% paraformaldehyde in 1xPBS for 4 hr at room temperature. Retinae were dissected and processed for flat mount. Specimens were permeabilized and blocked in 0.5% Triton X-100/5% normal goat serum for 12 hr at 4 °C before incubation with primary antibodies for 36-48 hr followed by multiple washes with 1xPBS and incubation for 2 hr with anti-goat, anti-rabbit or anti-mouse AlexaFluor-488, AlexaFluor-594 or Cy5-conjugated secondary antibodies. Hoechst (Invitrogen, CA) was used to counter-stain nuclei. Retina whole mounts were placed on glass slides with retinal ganglion cell side up with Fluoromount-G (Southern Biotech, Birmingham, AL). Images were acquired with a Nikon C1^+^ laser-scanning confocal microscope coupled with a LU4A4 launching base of four solid state diode lasers (407 nm/100 milliwatts, 488 nm/50 milliwatts, 561 nm/50 milliwatts, and 640 nm/40 milliwatts) and controlled by Nikon EZC1.3.10 software (version 6.4). Pan views of YFP^+^-RGNs from the marginal to optic nerve head regions of the retina were generated by stitching overlapping field of views of these neurons with Photoshop CS4 or Nikon Elements.

### Morphometric analyses

Quantitation of the numbers of YFP^+^Thy1^+^ and Brn3^+^-RGNs of control (+/+) and −/− mice was performed on images taken from retinal flat mounts (4 image fields of 127μm^2^ for both central and peripheral region, 3 mice/genotype). Axon diameters and g-ratios were measured and calculated from transmission electron microscopic images of cross-sections of optic nerves. The g-ratio of was calculated by the ratio between the averages of the maximal and minimal diameters of axon and myelinated fiber using NIKON element software AR (Ver. 4.0). A minimum of 100 randomly chosen axons per image field from at least 3 non-overlapping images per mouse were used. Quantitative analysis of the number and size of F4/80^+^-microglia in the ganglion cell layer of the retina was performed by counting and measuring positive immune fluorescence signals in the total image fields of 3 mm^2^ (approximately a quarter of whole retina was scanned, 3-4 mice/genotype). Surface areas of collapsed confocal stacks of microglia from retinal flat mounts were determined by image thresholding (image segmentation) of calibrated regions of interest (ROI) with Metamorph v7.0 (Molecular Devices).

### Immunoblotting

After mice were sacrificed by cervical dislocation and decapitation, the retinae were carefully dissected and the optic nerves were cut out just before optic chiasma (~5 mm). Tissues were snap frozen and placed on dry ice upon collection and stored at −80°C. Tissue homogenates were prepared as described previously with minor modifications [33,48]. Briefly, retinae were homogenized in radioimmune precipitation assay (RIPA) buffer with zirconium oxide beads (Next Advance, Averill Park, NY, ZROB05) and a Bullet blender (Next Advance, BBX24) at 8,000 rpm for 3 mins, whereas optic nerves were homogenized with stainless beads (Next Advance, SSB02) at 9,000 rpm for 2 mins with a Bullet blender^®^ (Next Advance, BBX24). Protein concentrations of tissue homogenates were measured by the BCA method using BSA as the standard (Pierce). Equal amounts of homogenates (60 μg of retina homogenates or 30 μg of optic nerve homogenates) were loaded and resolved in 7.5 % SDS-PAGE Hoefer or 4-15% gradient Criterion gels (BioRad, Hercules, CA). Blots were also reprobed for hsc70, whose protein levels were unchanged between genotypes, for normalization and quantification of proteins. Unsaturated band intensities were quantified by densitometry with Metamorph v7.0 (Molecular Devices, San Jose, CA), and integrated density values (idv) of bands were normalized to the background and idv of hsc70 as described previously [33,48].

### Biochemical assays

Retinae and optic nerves were collected immediately after the mice were euthanized, snap frozen and placed on dry ice, and stored at −80 °C in a freezer. NP-40 extracts were prepared with Bullet blender (Next Advance, BBX24). Free fatty acids (FFA) were measured with the FFA Quantification kit as described [34,55] and as per manufacturer’s instructions (Biovision, Mountain View, CA). A colorimetric assay kit for acetylcholinesterase activity was used as directed per manufacturer (Biovision) and previously described [34]. Data were normalized against protein amounts in NP40-solubilized tissue extracts used for each assay. Protein concentrations of NP40-solubilized extracts were determined by the Bradford assay (BioRad).

### Immunoprecipitation assays

Fresh retinal extracts were solubilized in Nonidet P-40 buffer using Bullet Blender BBX24 (Next Advance Inc.) in the presence of 0.5mm zirconium oxide beads (Next Advance Inc., Averill Park, NY). 1.2 mg of retinal extracts were pre-cleared with 2μg of non-immunized IgG (Stressgen, San Diego, CA) and incubated with 50μl of 50% protein A/G bead slurry (Santa Cruz Biotechnology) at 4°C for 1 hour. The supernatants were incubated with 2μg of mouse anti-HA antibody (Abcam, Cambridge, MA) at 4°C for 12 hour. Coimmunoprecipitate complexes were resolved by SDS-PAGE in 4-15% gradient Criterion gels (BioRad) and immunoblotted with mouse anti-Pttg1 antibody (Abcam) as described [33,48].

### Total RNA isolation and qRT-PCR

Total RNA was isolated using TRIzol^®^ (Invitrogen) following manufacturers’ guide. RNA was reverse transcribed using SuperScript II First-Strand Synthesis System (Invitrogen). Quantitation of mRNA level with gene-specific primers was carried out with cDNA equivalent to 10 ng of total RNA, SYBR Green PCR Master Mix and ECO Real-Time PCR System (Illumina Inc., San Diego, CA). The data were analyzed using Eco Real-Time PCR System Software version 4.0 (Illumina Inc.). The relative amount of transcripts was calculated by the ΔΔCT method. Data were normalized to GAPDH (*n*=3–4). Data were analyzed by Student’s *t*-test and a *p*-value≤ 0.05 was considered significant.

### Deep RNA sequencing by RNA-Seq

The dissections of optic and sciatic nerves were carried out concurrently from the same mice as described [34]. In this this study, total RNA from optic nerves was isolated and subjected to deep RNA sequencing exactly as previously described.[34] Briefly, optic nerves were collected and incubated in RNA*later* (Ambion/Thermo Fisher Scientific, Waltham, MA) and snap frozen in liquid nitrogen. Samples were submitted to Otogenetics Corporation (Norcross, GA USA) for RNA-Seq. The Agilent Bioanalyzer or Tapestation and OD260/280 was used to assess the purity and integrity of total RNA. 1-2 μg of cDNA was generated with the Clontech SMARTer cDNA kit (catalog# 634925; Clontech Laboratories, Inc., Mountain View, CA), fragmented with Covaris (Covaris, Inc., Woburn, MA) or Bioruptor (Diagenode, Inc., Denville, NJ), profiled with Agilent Bioanalyzer or Tapestation, and submitted to Illumina library preparation using NEBNext reagents (catalog# E6040; New England Biolabs, Ipswich, MA). The Agilent Bioanalyzer or Tapestation was used to determine the quality, quantity and the size distribution of the Illumina libraries and the libraries were submitted for Illumina HiSeq2000 or HiSeq2500 sequencing. Paired-end 100 nucleotide reads were generated from RNAseq with a sequence depth between 45-70 million seq reads and checked for data quality using FASTQC (Babraham Institute, Cambridge, UK). Data were analyzed with DNAnexus (DNAnexus, Inc, Mountain View, CA) or the platform provided by the Center for Biotechnology and Computational Biology (University of Maryland, College Park, MD) [56]. Levels of individual transcripts were expressed as fragments per kilobase of exon per million fragments mapped (FPKM) and were obtained using Cufflinks. A *q*-value less than 0.05 was considered as statistically significant.

### Statistics

Mice were randomly assorted and experiments were performed blind until data analysis. Samples sizes/independent biological replicates were collected (power > 0.8) and these were comparable with other studies using the same mouse lines and genotypes [32,34,48]. Twoway repeated measure analyses of variance were used to analyze luminance-response functions for measures of VEP amplitude and timing. g-ratios and axonal diameters between groups were assessed with a *t*-test of difference between means using generalized estimating equations (GEE) to account for multiple nerves per mouse. The difference between groups adjusting for axonal diameter was assessed using generalized estimating equations with terms for group, axonal diameter and their interaction (SAS, Cary, NC). The Mann-Whitney test rank-sum test was used to examine areas of perikarya of RGNs. For all other assays, Student’s *t*-test for two group comparisons was used. Data are reported as average values ± SD, except otherwise specified. Differences among the groups were considered statistically significant when *p*-value ≤ 0.05.

## Results

### Mice with targeted Ranbp2 in Thy1^+^-retinal ganglion neurons (RGNs)

We used the mouse line, <u>s</u>ingle-neuron <u>l</u>abeling with <u>i</u>nducible <u>C</u>re-mediated <u>k</u>nock-out <u>H</u> transgenic line (*SLICK-H*), which expresses the yellow fluorescent protein (YFP) and tamoxifen-inducible Cre recombinase (CreER^T2^) under the control of neural Thy1 promoter in Thy1^+^-motoneurons and RGNs [47,57–59], to cross with another line harboring a *Ranbp2* floxed gene (*Ranbp2^flox/flox^*) [34,49]. This cross generated the mouse line, *SLICK-H::Ranbp2^flox/flox^*, which lacks *Ranbp2* in motoneurons and RGNs after tamoxifen administration [34] (Fig 1A) (unless otherwise noted, hereafter *SLICK-H::Ranbp2^flox/flox^* (−/−) are mice that underwent tamoxifen treatment). We have previously shown that *SLICK-H::Ranbp2^flox/flox^* mice with loss of *Ranbp2* in Thy1^+^-motoneurons develop ALS-like traits that culminate with the death of mice 10.5 days after a 5-day regimen of tamoxifen [34]. However, the effects of loss of Ranbp2 in Thy1^+^-RGNs are unknown and this was the focus of this study. Like in motoneurons [34], the retina of *SLICK-H::Ranbp2^flox/flox^* produce a recombinant transcript with a deletion of exon 2, fusion of exons 1 and 3, and premature stop codon in exon 3 as soon as day 0 after a 5-day regimen of tamoxifen (Fig 1B). The recombinant *Ranbp2* transcript has the translation potential of only the first 31 residues of the 3053 residues that comprise Ranbp2. Fundus fluorescence imaging of eyes of live wild-type and *SLICK-H::Ranbp2^flox/flox^* mice treated with tamoxifen showed no overt fundoscopic differences of YFP^+^-RGNs and fasciculation of their axons throughout the retina between genotypes at ten days after the last dose of tamoxifen (d10) (Fig 1C). In contrast to wild-type mice, confocal microscopy of retinal flatmounts showed that YFP^+^-RGNs lacked Ranbp2 at the nuclear rim at d0 (Fig 1D). Akin to motoneurons and other studies of other cell types [34,49], the absence of Ranbp2 did not affect the localization of other nucleoporins at the nuclear rim of YFP^+^-RGNs (e.g., Nup153 and Nup62) (Fig 1E). In addition, loss of Ranbp2 did not cause the degeneration of YFP^+^-and Brn3^+^-RGNs across the retina (Figs 1F and G).

**Figure 1.**
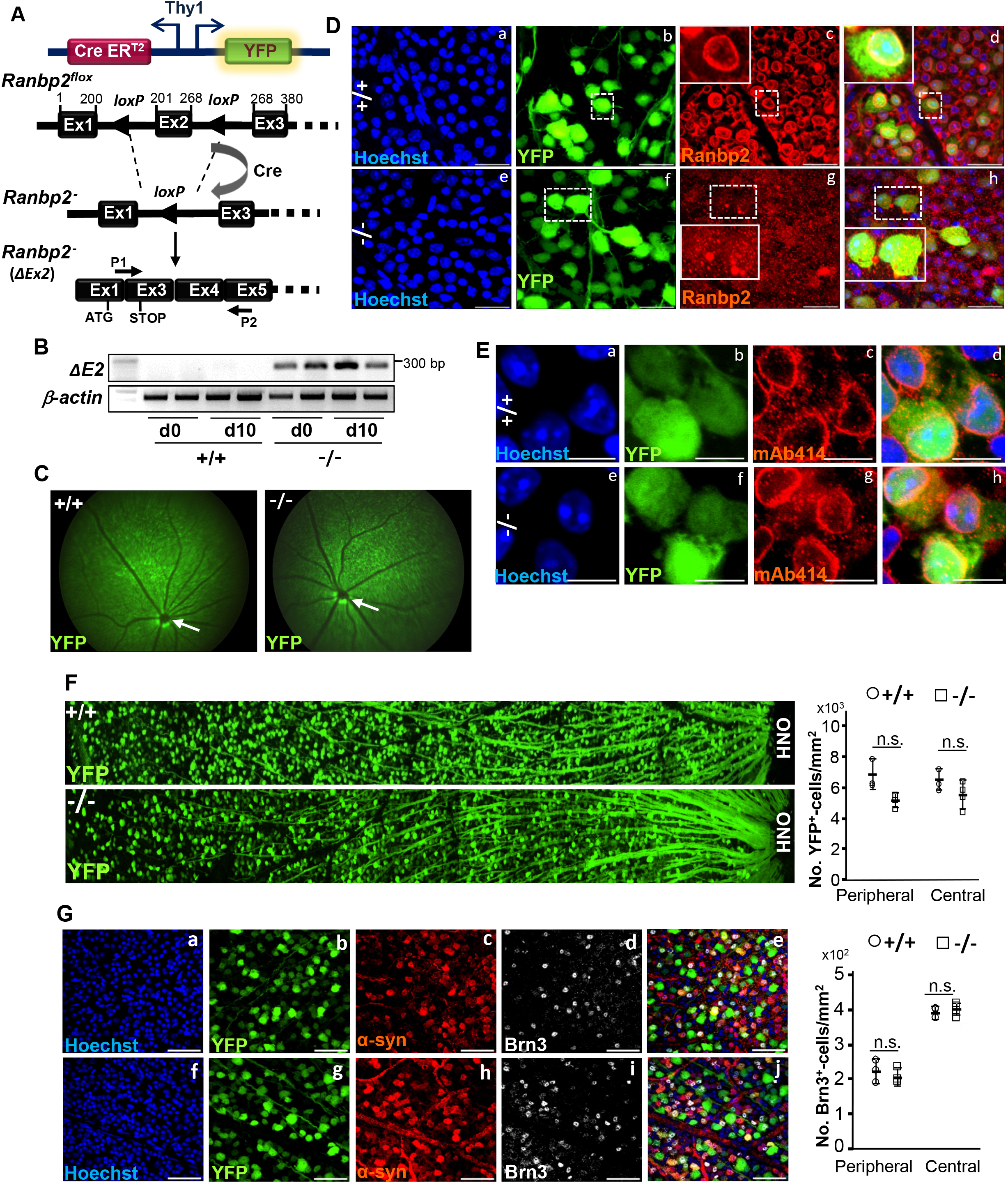
Genetic ablation of *Ranbp2* in YFP ^+^Thy1^+^-retinal ganglion neurons (RGNs). **A.** The single-neuron labeling with inducible Cre-mediated knock-out H transgenic line (SLICK-H) harbors two oppositely oriented *Thy1* promoters that control the co-expression of the yellow fluorescent protein (YFP) and tamoxifen-inducible Cre recombinase (CreER^T2^). This line was used to excise exon 2 (ΔEx2) from a floxed *Ranbp2* gene in mice and generate an out-of-frame exon 3 after splicing of exons 1 and 3. P1 is a hybrid primer against the junction produced by the splicing (fusion) of exons 1 and 3 of the recombinant transcript. P2 is a primer specific for exon 5. **B.** Expression of *Ranbp2* mRNA without exon 2 is detected in the retina by RT-PCR with P1 and P2 primers at the end of a daily 5-day regimen of tamoxifen administration, day 0 (d0), and ten days (d10) after the last dose of tamoxifen. **C**. Live fluorescence retinal fundoscopy of dilated pupils of mice showed no overt differences between +/+ and −/− mice at 10 days (d10) after the last dose of tamoxifen. Arrows denote optic nerve head. **D**. Confocal images of retinal flat mounts (RGNs facing up) show that compared to +/+ mice (a-d), YFP^+^-RGNs of −/− mice lack Ranbp2 at the nuclear envelope at d10 (e-h). Inset picture is an enlarged view of dashed-line box. **E**. Confocal images of retinal flat mounts (ganglion neurons facing up) show that the nucleoporins 153 and 62 (Nup153/Nup62) at the nuclear envelope and detected by mAb414 are not affected in YFP^+^-RGNs between +/+ (a-d) and −/− mice (e-h) at d10. **F**. Low-power confocal images from the optic nerve head (ONH) to the marginal border of the retina (left) and morphometric analyses (right) of retinal flat mounts (ganglion neurons facing up) show that there are no differences of YFP^+^-RGNs between +/+ and −/− mice at d10. Student’s *t*-test, *n*=4 mice/genotype; data are expressed as mean ± sd. **G**. Confocal images (right) and morphometric analyses (left) of retinal flat mounts (ganglion neurons facing up) show that there are no differences Brn3^+^-RGNs between +/+ and −/− mice at d10. There are also no apparent differences in -Syn^+^-RGNs between +/+ and −/− mice at d10. Student’s *t*-test, *n*=4 mice/genotype; data are expressed as mean ± sd. +/+, *SLICK-H::Ranbp2^+/+^*; −/−, *SLICK-H::Ranbp2^flox/flox^*; mAb414, monoclonal Ab414 against Ranbp2(Nup358)/Nup153/Nup62; scale bars=25 μm (D), 10 μm (E), and 50 μm (G); d0 and d10 are days 0 and 10 after the last dose of a daily 5-day regimen of tamoxifen administration, respectively; α-syn, α-synuclein; Brn3, Brn3a/b/c POU-domain transcription factors; OPN, optic nerve head; n.s., not significant.

### *SLICK-H::Ranbp2^flox/flox^* mice have a delay of implicit times of visual evoked potential (VEP)

We previously found that the sciatic nerves of *SLICK-H::Ranbp2^flox/flox^* present lower nerve conduction velocity and hind limb paralysis at day 9 after the last dose of tamoxifen [34]. Functional disturbances in the visual pathway from the RGNs to the primary visual cortex have been reported in ALS patients [42]. To ascertain how loss of Ranbp2 in YFP^+^-RGNs affected the transmission of luminance stimuli to the visual cortex, we measured the amplitude and latency (implicit times) of VEPs as a function of increasing flash luminance in wild-type and *SLICK-H:: Ranbp2^flox/flox^* mice before and after treatment with tamoxifen [60,61]. As shown in Fig 2A, there were no overt differences in VEP waveforms between untreated and tamoxifen-treated wild type mice, whereas there were noticeable changes in the implicit time of the dominant negative wave component (N1) of the VEP waveforms to strobe flash stimuli between untreated and tamoxifen-treated *SLICK-H::Ranbp2^flox/flox^* mice at day 5 (d5) after the last dose of tamoxifen [50]. Luminance-response functions for the N1 implicit time and amplitude of VEPs showed that there was a significant delay in the N1 implicit time of tamoxifen-treated *SLICK-H:: Ranbp2^flox/flox^* compared to wild-type mice (p<0.001; Fig 2B), while the amplitudes of VEPs did not differ between genotypes (Fig 2C).

**Figure 2.**
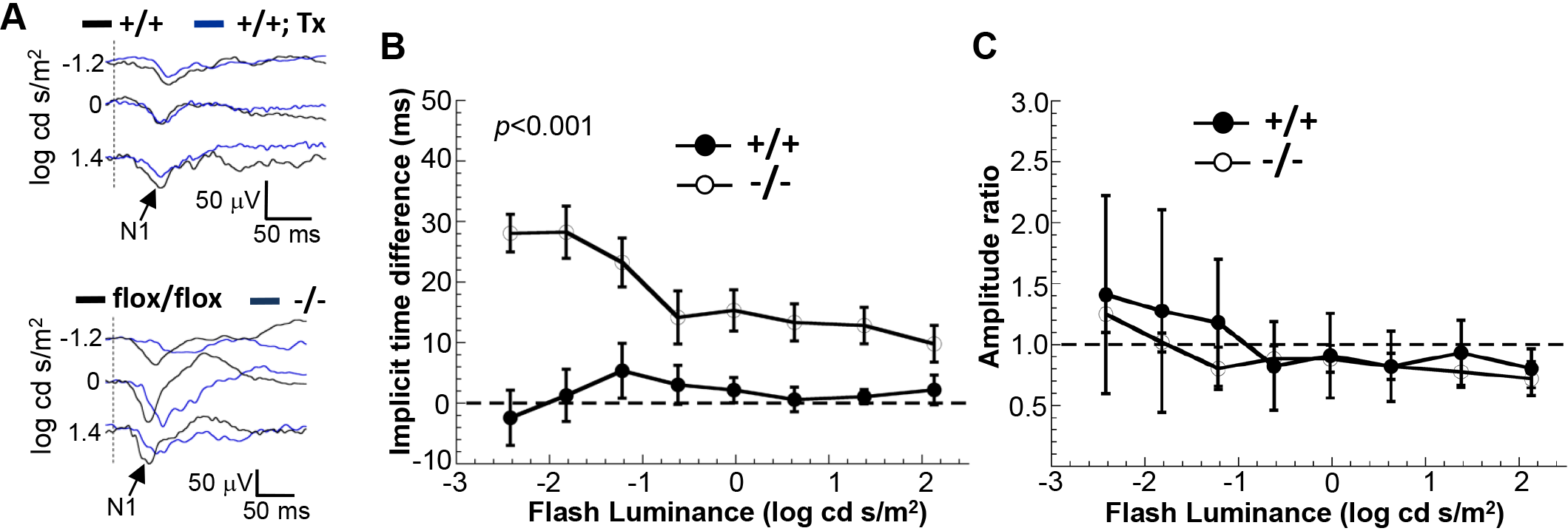
Electrophysiological deficits in the visual pathway of *SLICK-H::Ranbp2^flox/flox^*mice. **A**. Visual-evoked potentials (VEPs) recorded to a series of strobe flash stimuli from representative tamoxifen (Tx) untreated (black and baseline tracings) and treated (blue tracings) +/+ (top panel) and *flox/flox* mice (bottom panel). N1, the major negative component of the mouse VEP, is indicated by arrows in the tracings. In comparison to baseline, the VEP obtained following tamoxifen is slower in −/− than +/+ and *flox/flox* mouse. The vertical dashed lines indicate the strobe flash stimulus presentation. **B.** Average (± sem) ratio of VEP implicit times measured at baseline and five days (d5) following a daily 5-day regimen of tamoxifen administration for +/+ and −/− mice. In comparison to +/+ controls, two-way repeated measures ANOVAs indicate that VEP implicit times were significantly delayed in −/− mice (*p*<0.001). **C.** Average (± sem) ratio of VEP amplitude measured at baseline and 5 days (d5) following a daily 5-day regimen of tamoxifen administration for +/+ and −/− mice. Two-way repeated measures ANOVAs indicate that VEP amplitude was not significantly affected between genotypes (*p*>0.05). *n*=6 mice/genotype. +/+, *SLICK-H::Ranbp2^+/+^*; −/−, *SLICK-H::Ranbp2^flox/flox^*; *flox/flox,* tamoxifen-untreated *SLICK-H::Ranbp2^flox/flox^*.

### RGNs and optic nerves of *SLICK-H::Ranbp2^flox/flox^* mice develop morphometric abnormalities

To gain further insights into the bases of the impairment of VEPs in *SLICK-H::Ranbp2^flox/flox^* mice, we compared morphometric analyses of myelinated axons and cell bodies of RGNs between genotypes. Examination of transmission electron microscopy images of RGNs between untreated and tamoxifen-treated *SLICK-H::Ranbp2^flox/flox^* mice found no overt differences in the ultrastructural morphology of the somata of these neurons at d10 (Fig 3A). However, morphometric analyses showed a significant increase of the mean perikarya area (hypertrophy) of YFP^+^-RGNs of *SLICK-H::Ranbp2^flox/flox^* mice compared to *SLICK-H::Ranbp2^+/+^* by d10 (*p*=0.00005) (Fig. 3B). Next, we examined ultrathin sections of the optic nerve at d10 and found that there were also no overt and gross differences in morphology of myelinated axons of the optic between genotypes (Fig. 3C).

**Figure 3.**
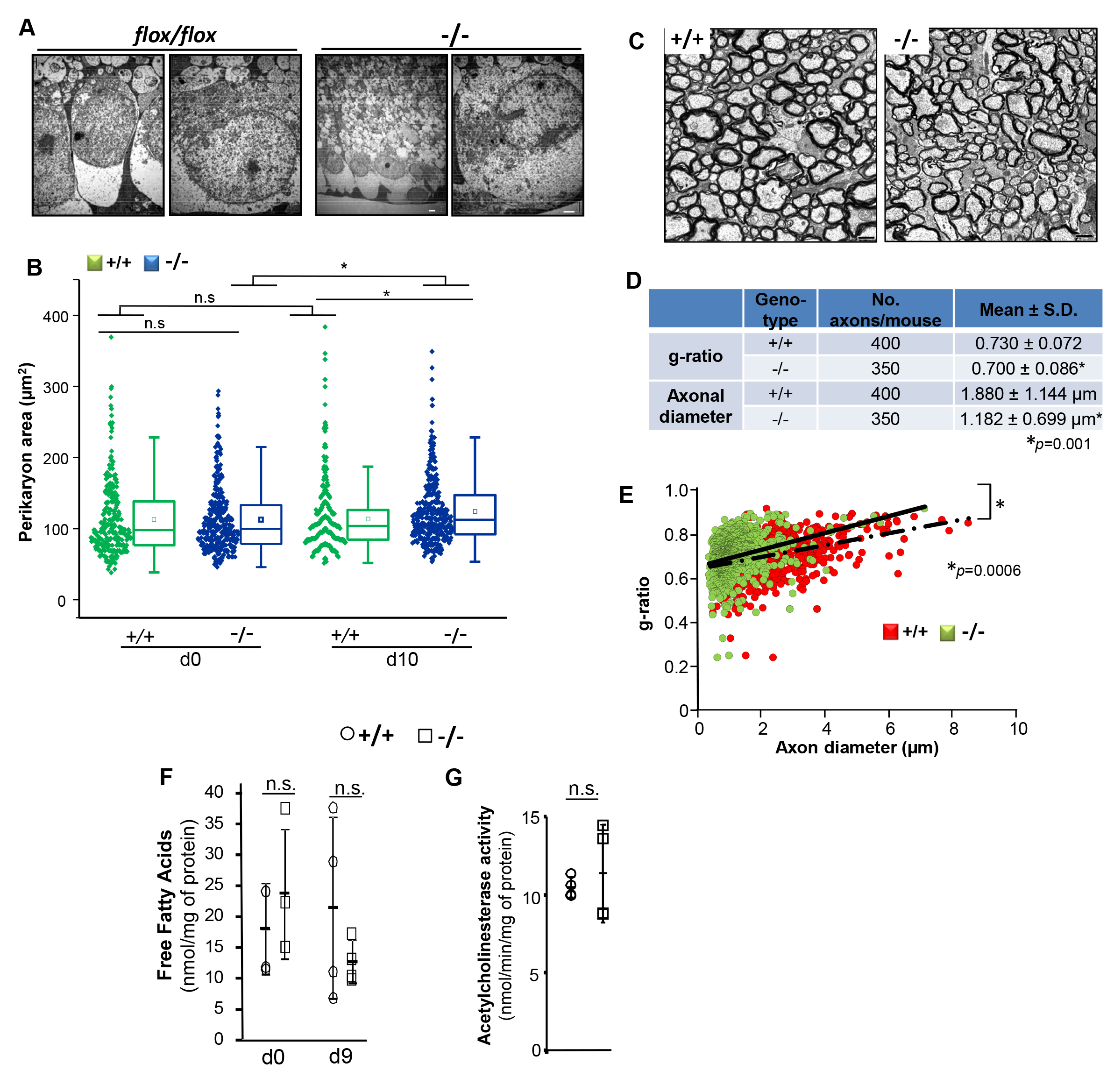
Morphometric and lipid profile changes of retinal ganglion neurons (RGNs) of *SLICK-H::Ranbp2^flox/flox^* mice. **A.** Representative low (left) and high power (right) transmission electron micrographs of RGNs of the retina of *flox/flox* (without tamoxifen treatment) and −/− mice at day 10 (d10) post-tamoxifen administration. No overt ultrastructural changes in somata of RGNs were observed between tamoxifen-treated and untreated mice. **B**. Dot-box plots of nonparametric analyses of perikarya (soma) area of YFP^+^-RGNs. The perikarya of YFP^+^-RGNs of – /− mice develop hypertrophy. Box-plot edges denote the 25th and 75th percentiles of the data, and the centerline denotes the median value. *n*=291(+/+, d0), 340 (+/+, d10), 332 (−/−, d0), 335 (−/−, d10); \**p*≤0.00005, Mann-Whitney test, *n*=4 mice/genotype. **C**. Representative cross-sections of transmission electron micrographs of myelinated optic nerves of +/+ and −/− mice at d10. **D**. Measurements of g-ratios and axonal diameters of optic nerves showed significant decreases between genotypes. Data are expressed as mean ± sd., \**p*<0.001, *t*-tests with GEE, *n*=4 mice/genotype. **E**. Scatter plots between g-ratios and axonal diameter of fibers of optic nerves. Slopes of +/+ and −/− mice are 0.0269 ± 0.033 and 0.0385 ± 0.0006, respectively; \**p*<0.0006 (interaction term), *t*-tests with GEE, *n*=4 mice/genotype. Slope values are expressed as mean ± 40 s.e.m. Slopes diverged with the increase of axon caliber. **F.** The levels of free fatty acids are unchanged between gentotypes in the optic nerve at d9 (Student’s *t*-test, *n*=4 mice/genotype). Data are expressed as mean ± sd. **G.** The levels of acetylcholinesterase activity are unaltered between genotypes in the visual cortex at d9 (Student’s *t*-test, *n*=4 mice/genotype). Scale bars= 2 μm (**A, C**). n.s., non-significant; −/−, *SLICK-H::Ranbp2^flox/flox^*; +/+, *SLICK-H::Ranbp2^+/+^*; d0, d9 and d10 are days 0, 9 and 10 post-tamoxifen administration, respectively.

To uncover potential changes in myelination and axonal diameters of the fibers of the optic nerve, we measured the g-ratio and axonal diameters. The *g*-ratio quantifies changes in the ratio between the diameter of the inner axon alone and that of the axon with myelin sheath. We found that by d10 the *g*-ratio were slightly but significantly diminished between *SLICK-H:: Ranbp2^flox/flox^* and *SLICK-H::Ranbp2^+/+^* (*p*=0.001), whereas the diameters of axons of the optic nerve were strongly decreased by ~40% in *SLICK-H::Ranbp2^flox/flox^* compared *SLICK-H:: Ranbp2^+/+^* (*p*=0.001) (Fig. 3D). A decrease in g-ratio typically indicates hypermyelination, but a decrease of axonal caliber alone also contributes to a decline in *g*-ratio.[62] Hence, we built scatter plots of g-ratios of axons of optic nerve as a function of the axonal diameter to examine further relationships between g-ratio and axon caliber of the optic nerve (Fig. 3E). This analysis showed that compared to *SLICK-H::Ranbp2^+/+^*, there was a significant leftward shift in the scatter plot of *SLICK-H::Ranbp2^flox/flox^* mice indicating that large caliber axons were affected by a strong decrease in axon diameter (Fig. 3E). This observation was also supported by the examination of the slopes of the regression lines (0.0269 ± 0.033 *vs* 0.0385 ± 0.0006) that diverged with the increase of axon caliber and that were significantly different between genotypes (*p*=0.0006) (Fig. 3E). Likewise, the slope for each genotype and optic nerve was significant (*p*<0.0001). Collectively, these data indicate that large caliber axons of *SLICK-H:: Ranbp2^flox/flox^* mice exhibit an increased vulnerability to degeneration as reflected by a selective and major decrease of axonal diameter of high caliber fibers and that these effects were accompanied by the hypertrophy of the soma of YFP^+^-RGNs of *SLICK-H::Ranbp2^flox/flox^* mice.

We have previously found that the levels of free fatty acids (FFA) were decreased in the sciatic nerve of *SLICK-H::Ranbp2^flox/flox^* mice [34]. Hence, we examined also the levels of free fatty acids in the optic nerve between genotypes. We found that the levels of FFA (octanoate and longer fatty acids) remained unchanged between genotypes (Fig. 3F). Finally, we examined the levels of acetylcholinesterase (AChE) activity in the visual cortex, since cholinergic synapses of motoneurons may be affected during the course of some forms of ALS and acetylcholine (ACh) modulates the processing of visual information in the visual cortex (V1)[63]. However, and like in motoneurons [34], the activity of AChE was similar in the visual cortex between genotypes at d9 (Fig. 3G).

### RGNs of *SLICK-H::Ranbp2^flox/flox^* mice present impairments in the nuclear-cytoplasmic distribution of direct and accessory partners of Ranbp2

Ranbp2 is a large modular protein, whose domains interact with specific partners (Fig. 4A). In particular, the Ran-binding domains (RBDs) of Ranbp2 associate with Ran-GTP and the accessory nuclear import receptor, importin-β[20,21], whereas the zinc-finger-rich domain (ZnF) is a docking site for the nuclear export receptor, exportin-1 (Fig. 4A) [26]. We have shown previously that the nucleocytoplasmic distributions of these receptors, Ran GTPase and accessory substrates of Ranbp2 become mislocalized between the nuclear and cytosolic compartments after loss of Ranbp2 or selective activities of its domains in motoneurons and other cell types [32–34]. These nuclear export and import receptors may also control the nuclear shuttling of NR2F2, an orphan nuclear receptor that interacts with Ranbp2 in the retina [54]. In addition, HDAC4 is a substrate of Ranbp2 [64] and Ranbp2 controls the nucleocytoplasmic distribution of HDAC4 in motoneurons [34] and its proteostasis via the ubiquitin proteasome system (UPS) [65]. Hence, we examined the effect of loss of Ranbp2 in the nucleocytoplasmic localizations of these direct and accessory partners of Ranbp2 in RGNs (Figs. 4B-D). We found that importin-β and exportin-1 were localized at the nuclear rim and intranuclear compartment of YFP^+^-RGNs of wild-type mice, whereas there was strong accumulation of importin-β and exportin-1 in the cytoplasmic compartment of YFP^+^-RGNs of *SLICK-H::Ranbp2^flox/flox^* (Fig 4B). These effects were also accompanied by the general decrease of NR2F2^+^-nuclei of YFP^+^-RGNs of *SLICK-H::Ranbp2^flox/flox^* compared to wild type mice (Fig 4C). Like in motoneurons [34], HDAC4 was largely excluded from the nuclear compartment of YFP^+^-RGNs of wild type mice, but this subcellular compartmentalization was lost in *SLICK-H::Ranbp2^flox/flox^* (Fig 4C). Ran GTPase and its nucleotide-bound form, Ran-GTP, were found diffusely dispersed throughout the perikarya and nuclei of YFP^+^-RGNs of wild type mice (Fig 4D), but Ran-GTP was also found at the nuclear rim of YFP^+^-RGNs (Fig 4D, a”’). By contrast, Ran GTPase localization appeared redistributed mostly to the cytosolic compartment, and Ran-GTP localization in the nucleus and at the nuclear rim was largely lost, in YFP^+^-RGNs of *SLICK-H::Ranbp2^flox/flox^* (Fig 4D).

**Figure 4.**
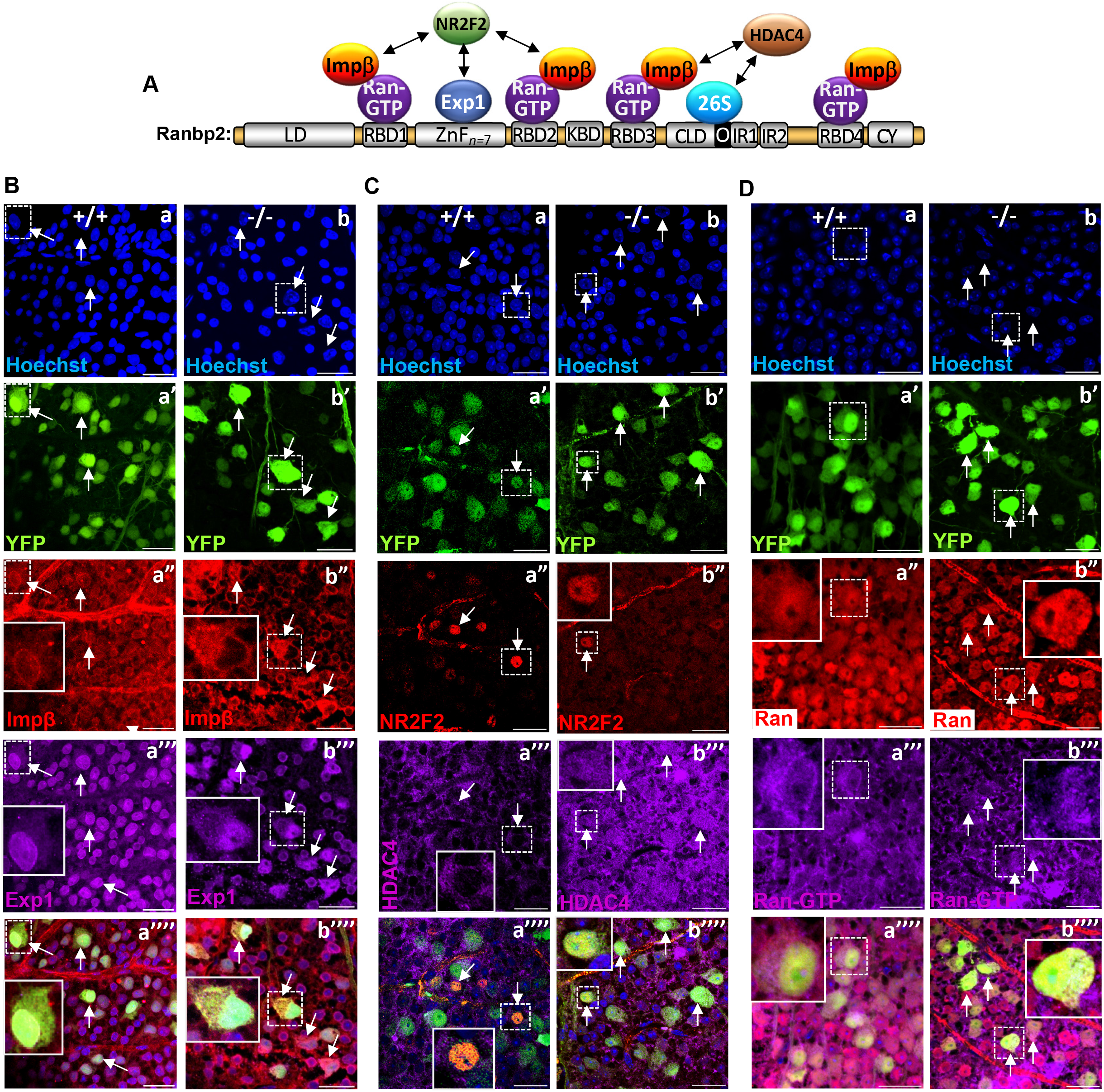
Disruption of subcellular localization of partners and substrates of Ranbp2 in retinal ganglion neurons (RGNs) of *SLICK-H::Ranbp2^flox/flox^* mice. **A.** Schematic diagram of the multimodular structure of Ranbp2 and its partners. **B-D.** Confocal images of RGNs of retinal flat mounts (ganglion neurons facing up) at d10 (day10) post-tamoxifen administration and coimmunostained with partners and substrates of Ranbp2. Inset pictures are enlarged views of dashed-line boxes. **(B)** Importin-β (a’’) and exportin-1 (a’’’) localize to the nuclear rim and nucleus of YFP^+^-RGNs of +/+ mice (arrows). In −/− mice, YFP^+^-RGNs have cytosolic accumulation of importin-β(b’’), and cytosolic and intranuclear accumulation of exportin-1 (b’’’) (arrows). **(C)** NR2F2 (a’’) localizes to nuclei, whereas HDAC4 (a’’’) is excluded from the nuclei compartment of YFP^+^-RGNs of +/+ mice. In −/− mice, there is loss of NR2F2^+^-RGNs (b’’) and of the subcellular partitioning of HDAC4 in YFP^+^-RGNs (b’’’). **(D)** Ran GTPase (a’’) and Ran-GTP (a’’’) localize diffusely throughout the cytosolic and nuclear compartments of YFP^+^-RGNs of +/+ mice and Ran-GTP localizes also to the nuclear rims of YFP^+^-RGNs (a’’’). In −/− mice, there is subcellular redistribution of Ran GTPase (b’’) and Ran-GTP (b’’’) into the cytosolic compartment and loss of Ran-GTP at the nuclear rims of YFP^+^-RGNs. Scale bars=25μm. LD, leucine-rich domain; RBD_n=1–4_, Ran GTPase-binding domains, n=1–4; ZnF_n =7_, zinc finger-rich domains; KBD, kinesin-1-binding domain; CLD, cyclophilin-like domain; IR1 and IR2, internal repeats 1 and 2, respectively; M, middle domain between IR1 and IR2; O, overlapping region between CLD and IR1; CY, cyclophilin domain; Exp1, exportin-1; impβ, importin-β; HDAC4, histone deacetylase 4; Ran, Ran GTPase, 26S, subunits of the 26S proteasome; NR2F2, nuclear receptor 2 factor 2; −/−, *SLICK-H::Ranbp2^flox/flox^*; +/+, *SLICK-H:: Ranbp2^+/+^*.

### Differential transcriptomics and gene expression analysis of optic nerves between wild-type and *SLICK-H::Ranbp2^flox/flox^*

Our prior differential transcriptome and gene expression studies of sciatic nerves between wild-type and *SLICK-H::Ranbp2^flox/flox^* mice uncovered the regulation of 25 transcripts by Ranbp2 including a set of chemokine receptors and ligands [34]. To gain insights into transcriptome changes that underpin the pathobiological phenotypes shared by and unique to sciatic and optic nerves after nucleocytoplasmic impairment caused by loss of *Ranbp2*, we performed differential and whole transcriptome analysis of optic nerves that are comprised largely by axons of RGNs and oligodendrocytes (myelin sheaths) by deep RNA sequencing (RNAseq) of 45-70 million total seq reads of *SLICK-H::Ranbp2^+/+^, SLICK-H::Ranbp2^flox/flox^* and *Tg*-*Ranbp2^RBD2/3*-HA^*::*SLICK-H::Ranbp2^flox/flox^* (Fig. 5A). The *Tg*-*Ranbp2^RBD2/3*-HA^* ::*SLICK-H:: Ranbp2^flox/flox^* expresses a BAC transgene with mutations in RBD2 and RBD3 of Ranbp2 in a *null Ranbp2* background that abolish their association to Ran-GTP and importin-β[32,34]. This line served as a control line, because expression of *Tg*-*Ranbp2^RBD2/3*-HA^* rescues the lethality and motor behaviors of *SLICK-H::Ranbp2^flox/flox^* and thereby serves to filter out compensatory transcriptional responses without pathophysiological relevanc [34,32].

**Figure 5.**
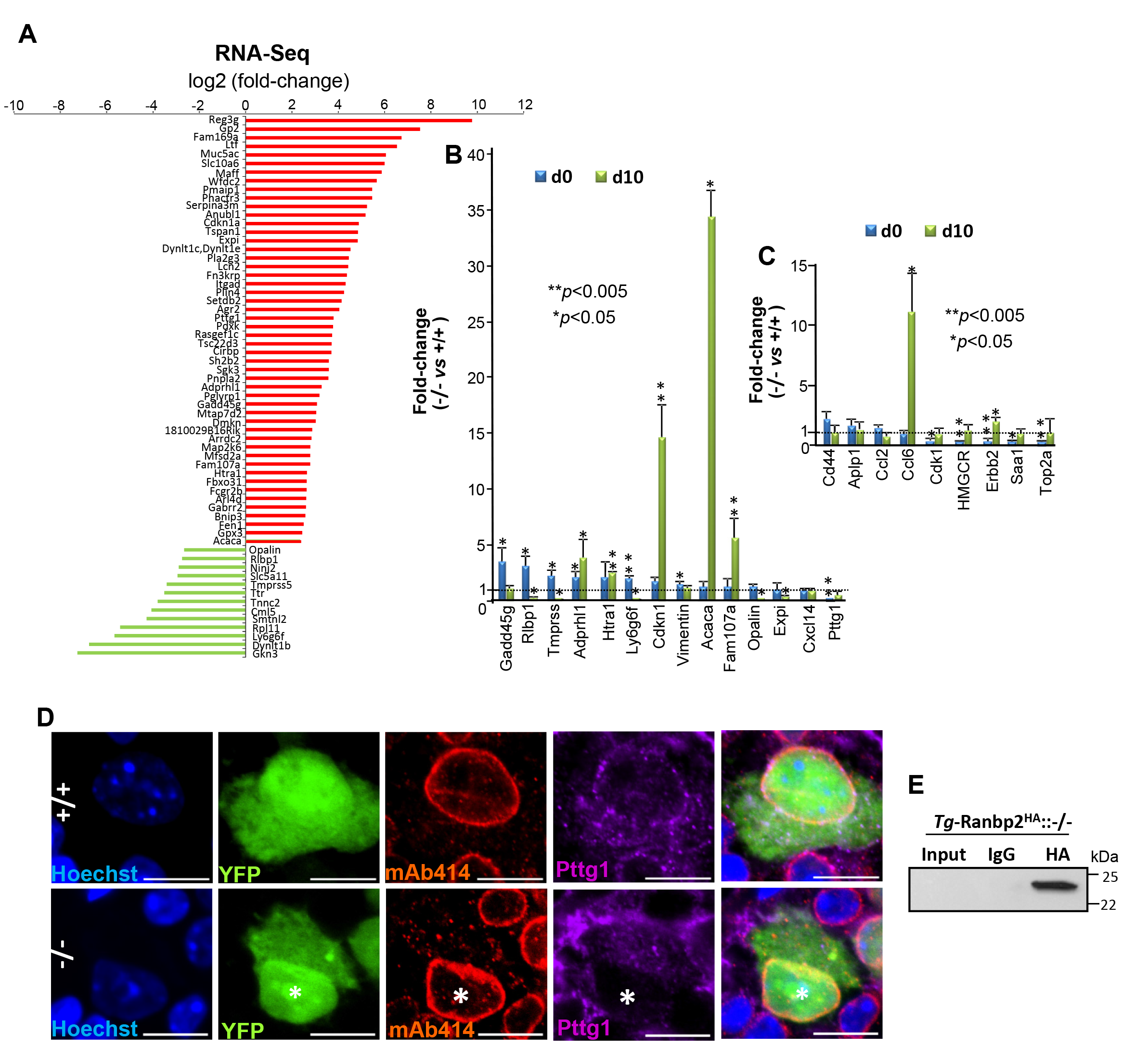
Differential transcriptome and gene expression analyses of optic nerve of retinal ganglion neurons (RGNs) upon loss of *Ranbp2*. **A.** Differential RNA-Seq-based wholetranscriptome analysis of optic nerves between *SLICK-H::Ranbp2^flox/flox^* (−/−), *SLICK-H:: Ranbp2^+/+^* (+/+) and *Tg^RBD2/3*-HA^::SLICK-H::Ranbp2^flox/flox^* at day 10 post-tamoxifen administration. Fifty transcripts were up-regulated (red) and thirteen transcripts were downregulated in −/− mice at d10. **B.** Validation of ranked RNA-Seq dataset by RT-qPCR and temporal and directional changes of expression levels of mRNAs between −/− and +/+ mice. Thirteen transcripts were validated to be up-regulated and *Pttg1* was validated to be downregulated by RT-qPCR in −/− mice at d0, d10 or both. *Acaca* (acetyl-CoA carboxylase alpha; also known as Acc1) and cdkn1 (cyclin-dependent kinase inhibitor 1) had the strongest up-regulation (~35 and 15-fold, respectively) at d10, whereas Pttg1 was down-regulated by ~25-fold at d0. Data are expressed as mean ± sd. Student’s *t*-test, *n*=3-4 mice/genotype. **C.** Temporal and directional changes of expression of mRNAs in the optic nerve that are known to be differentially expressed in sciatic nerve by loss of *Ranbp2*. Five genes (*Ccl6*, *Cdk1*, *HMGCR*, *erbb2*, *Saa1* and *Top2a*) were found also to be deregulated in the optic nerve at d0, d10, or both. *Ccl6* had the strongest change of expression (~11-fold increase). Data are expressed as mean ± sd. Student’s *t*-test, *n*=3-4 mice/genotype. **D.** Confocal images of retinal flat mounts (ganglion neurons facing up) co-immunostained for Ranbp2 (Nup358)/Nup153/Nup62 (mAb414) and Pttg1. Pttg1 localizes at the nuclear rim with Ranbp2 (Nup358)/Nup153/Nup62 in YFP^+^-RGNs of +/+ mice, while Pttg1 localization is lost at the nuclear rim of YFP^+^-RGNs (nuclei labeled with *) of −/− mice. **E**. HA-tagged Ranbp2 (*Tg*-Ranbp2^HA^) expressed in transgenic mice with a *null Ranbp2* background (−/−) co-immunoprecipitates Pttg1 from retinal extracts. Pttg1 is not detected in an overloaded aliquot of input extracts (first lane) owing to its very low abundance in retinal extracts. −/−, *SLICK-H::Ranbp2^flox/flox^*; +/+, *SLICK-H::Ranbp2^+/+^; Tg-Ranbp2^HA^::*−/−, *Tg-Ranbp2^HA^::Ranbp2^−/−^*; d0 and d10 are days 0 and 10 post-tamoxifen administration, respectively.

To identify transcripts differentially modulated between genotypes at d10, we applied a cutoff with a log^2^ fold-change (FC) ≥ │4│ and false discovery rate (FDR) <0.05 (*q*<0.05) for the whole transcriptome analysis. As shown in Fig 5A, the differential transcriptome analysis identified 50 and 13 transcripts that were differentially up-regulated or down-regulated, respectively, between optic nerves of wild-type and *SLICK-H::Ranbp2^flox/flox^* (Fig. 5A). Then, we independently validated the relative magnitude, direction and temporal changes of the RNAseq transcripts in optic nerves between genotypes at d0 and d10 by RT-PCR (Fig. 5B). Among these transcripts, quantitative RT-PCR analysis validated 13 transcripts that were differentially modulated between genotypes at d0, d10 or both (Fig. 5B). We found that the transcripts with the highest up-regulation by d10 were acetyl-CoA carboxylase 1 (Acc1; ~35-fold, *p* value < 0.05), cyclin-dependent kinase inhibitor 1A (Cdkn1a; ~15-fold, *p* value < 0.05) and Family with sequence similarity 107/Down-regulated in renal cell carcinoma 1 (Fam107a/Drr1; ~6-fold, *p* value < 0.05). Among the transcripts with the highest down-regulation were the mammalian-specific myelin and paranodal protein, opalin (~0.14-fold at d10, *p* value < 0.05), the pituitary tumor-transforming gene I/securin (Pttg1; ~ 0.04-fold at d0, *p* value < 0.05) and lymphocyte antigen 6 family member G6F (Ly6g6f; ~ 0.10-fold at d10, *p* value < 0.05) (Fig. 5B). Notably, none of the transcripts differently regulated by Ranbp2 in the optic nerve overlapped with those that we previously reported by the same exact parameters in the sciatic nerve of the same mice [34]. To confirm further the cell-type specific effects of Ranbp2 between RGNs and motoneurons, we rescreened by qRT-PCR the optic nerve for transcriptional changes found to be modulated by Ranbp2 in the sciatic nerve. We found only 6 transcripts with significant transcriptional changes between sciatic and optic nerves. However, and with the exception of chemokine (C-C motif) ligand 6 at d10 (Ccl6; ~11-fold, *p* value < 0.05), the magnitude of the transcriptional changes were minor and below the cut-off used for differential analysis (< 2-fold) (Fig. 5C).

Given that the anti-apoptotic protein, Pttg1/securin, is a nuclear shuttling factor [66], we used the transcriptome findings to pursue the validation of this protein as a substrate of Ranbp2. We examined first its subcellular localization with the nucleoporins, Ranbp2/Nup358, Nup153 and Nup62, at the nuclear pore complexes (NPCs) of YFP^+^-RGNs of wild type and *SLICK-H:: Ranbp2^flox/flox^*. As shown in Fig 5D, the YFP^+^-RGNs of wild-type mice showed prominent localization of Pttg1/securin with NPCs at the nuclear rim. In comparison, the nuclear rims of YFP^+^-RGNs of *SLICK-H::Ranbp2^flox/flox^* mice lacked any immunostaining of Pttg1/securin. To examine further the association of Pttg1/securin with Ranbp2, we performed coimmunoprecipitation assays with a transgenic line expressing a wild-type *Ranbp2* transgene tagged with HA in a *null* background [33] (Fig. 5E). We found that Ranbp2 coimmunoprecipitated Pttg1/securin even though the abundance of Pttg1/securin was extremely low and could not be detected in immunoblots overloaded with aliquots of retinal extracts (Fig. 5E). This outcome likely reflects the low abundance of Pttg1/securin in RGNs and/or the very low representation of RGNs in the retinal neurome.

### Loss of Ranbp2 in YFP^+^-ganglion neurons causes the paracrine activation of microglia

As described, differential expression analyses between genotypes uncovered that acetyl-CoA carboxylase 1 (Acc1) had the strongest up-regulation (~35-fold) in optic nerves of YFP^+^-RGNs of *SLICK-H::Ranbp2^flox/flox^* mice. Acc1 is a critical fatty acid biosynthetic, rate-limiting and cytosolic isozyme, which mediates the conversion of acetyl-CoA to malonyl-CoA, a critical carbon donor for the synthesis of long-chain fatty acids [67,68]. Acc1 is under tight posttranslational and allosteric regulation and represents a critical metabolic switch in response to micro-environmental cues [68]. In particular, Acc1 was found recently to control T cell immunity [69,70]. For example, suppression of Acc1 impairs the proliferation of CD8^+^ T-cells [69] and the formation of interleukin-17-secreting T-cells of T-helper 17 (T_H_17), while promoting the differentiation of anti-inflammatory and regulatory T-cells (T_reg_) [70]. In the central nervous system, T_reg_ harbor alternative regenerative capacity by promoting remyelination and oligodendrocyte differentiation [71].

We examined the effects of loss of Ranbp2 in YFP^+^-RGNs in neuroimmunity by analyzing the paracrine stimulation of resident macrophages of the CNS, the microglia. In this respect, we used whole mount retinae immunostained with steady-state markers of microglia, such as CD11b and CD45, to examine the expression of these markers and morphological changes in microglia evoked by loss of Ranbp2 in YFP^+^-RGNs. The expression levels of CD11b and CD45 have been used to differentiate resting from activated parenchymal microglia [72]. Comparisons of retinae of wild type and *SLICK-H::Ranbp2^flox/flox^* mice by en face confocal microscopy showed that the retinal ganglion cell layer of *SLICK-H::Ranbp2^flox/flox^* mice had stronger CD11b^+^ (Fig. 6A) and CD45^+^-immunostaining (Fig. 6B) of ramified microglia than wild type mice (see also Supplementary Fig. 1).

**Figure 6.**
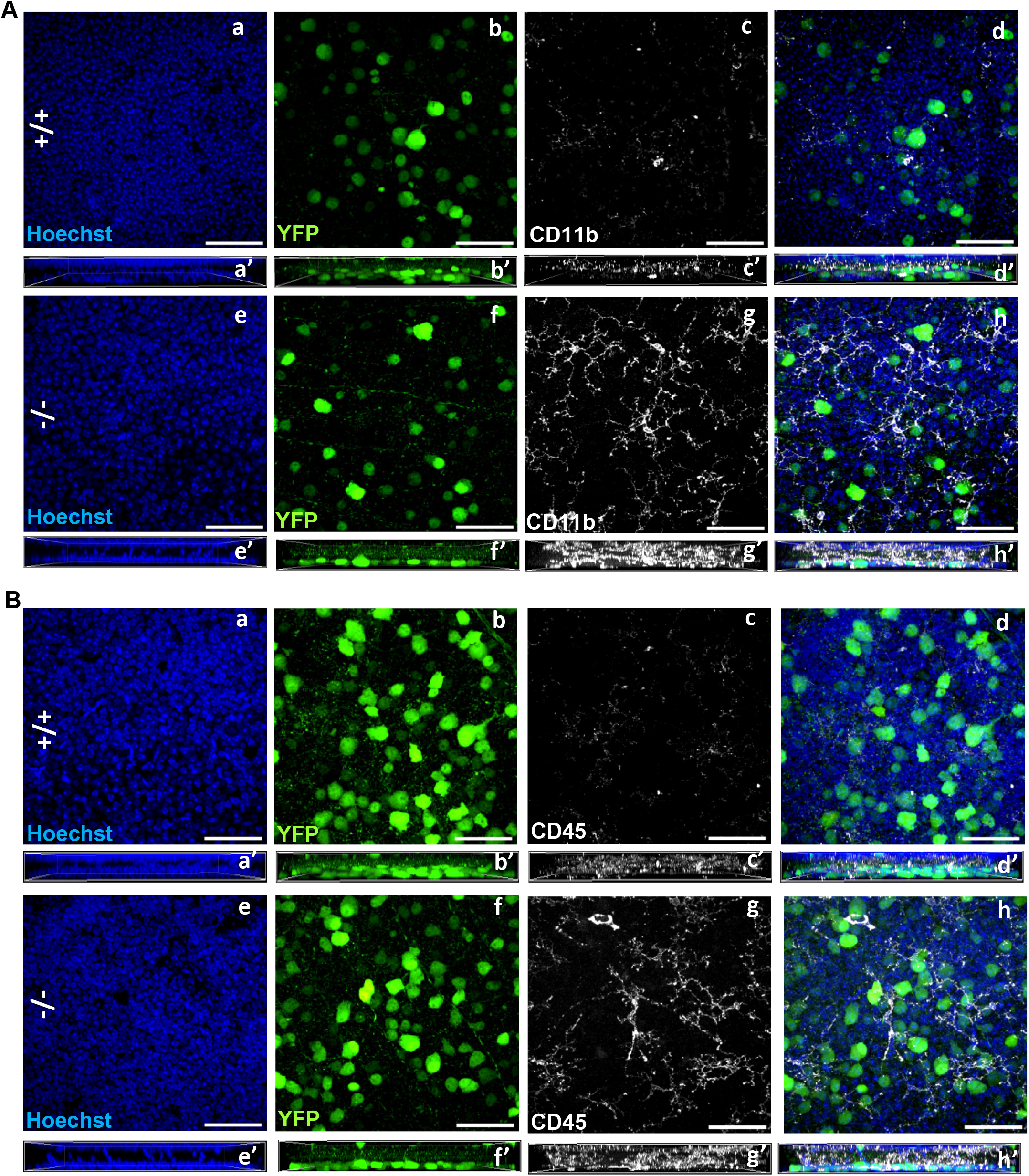
Activation of CD11b^+^ and CD45^+^-microglia in the ganglion cell layer of the retina by loss of *Ranbp2* in RGNs. Confocal images of retinal flat mounts (ganglion neurons facing up) of +/+ and −/− mice immunostained for CD11b (**A**) and Cd45 (**B**) at d10. Compared to +/+ (**A**, a-d), there is prominent activation of CD11b^+^-microglial cells, whose proliferative processes surround YFP^+^-RGNs of −/− mice (**A**, f-h). There is also prominent activation of CD45^+^-microglial cells, whose proliferative processes surround YFP^+^-RGNs of −/− mice (**B**, f-h). Figures a’-h’ in **A** and **B** represent lateral views of *z*-stacks of a-h and collapsed from a 25 μm thickstack of 13 images captured 2μm apart. Scale bars=50μm. d10, day 10 post-tamoxifen administration; −/−, *SLICK-H::Ranbp2^flox/flox^*; +/+, *SLICK-H::Ranbp2^+/+^*.

The transformation of microglia from resting to activated states is also known to be accompanied by changes in morphological phenotypes that is characterized by activated microglia losing their ramified or pseudopodial processes and gaining amoeboid appearances [72–74]. Hence, we used another marker for activated microglia, F4/80, to investigate morphological transformations in microglia caused by loss of Ranbp2 in YFP^+^-RGNs. As shown in Fig 7A, we observed that the RGN layer of wild type mice had intermingled F4/80^+^-cell bodies of various sizes, whereas scanning of the RGN layer of *SLICK-H::Ranbp2^flox/flox^* mice indicated that there was a higher abundance of F4/80^+^-cell bodies and that some of these cells had developed an amoeboid morphology. In light of the apparent non-uniform morphology of F4/80^+^-microglia between wild type and *SLICK-H::Ranbp2^flox/flox^* mice, we carried out morphometric analysis of F4/80^+^-microglia between genotypes. We found that *SLICK-H:: Ranbp2^flox/flox^* had an overall ~2-fold increase of F4/80^+^-microglia compared to wild type mice (Fig 7B; p<0.04). Then, we decomposed F4/80^+^-microglia intermingled between YFP^+^-RGNs based on their size. We found changes in the size of a subset of F4/80^+^-microglia between genotypes. In comparison to wild-type mice, *SLICK-H::Ranbp2^flox/flox^* mice had a significant ~2 (p<0.02) and ~4-fold (p<0.01) increase of amoeboid-like F4/80^+^-microglia with sizes of ~50 and ~100 μm^2^ and this increase was accompanied by a significant ~3-fold (p<0.01) decrease of F4/80^+^-microglia with extended processes like lamellipodial/pseudopodial and sizes of ~400 μm^2^ (Fig 7C).

**Figure 7.**
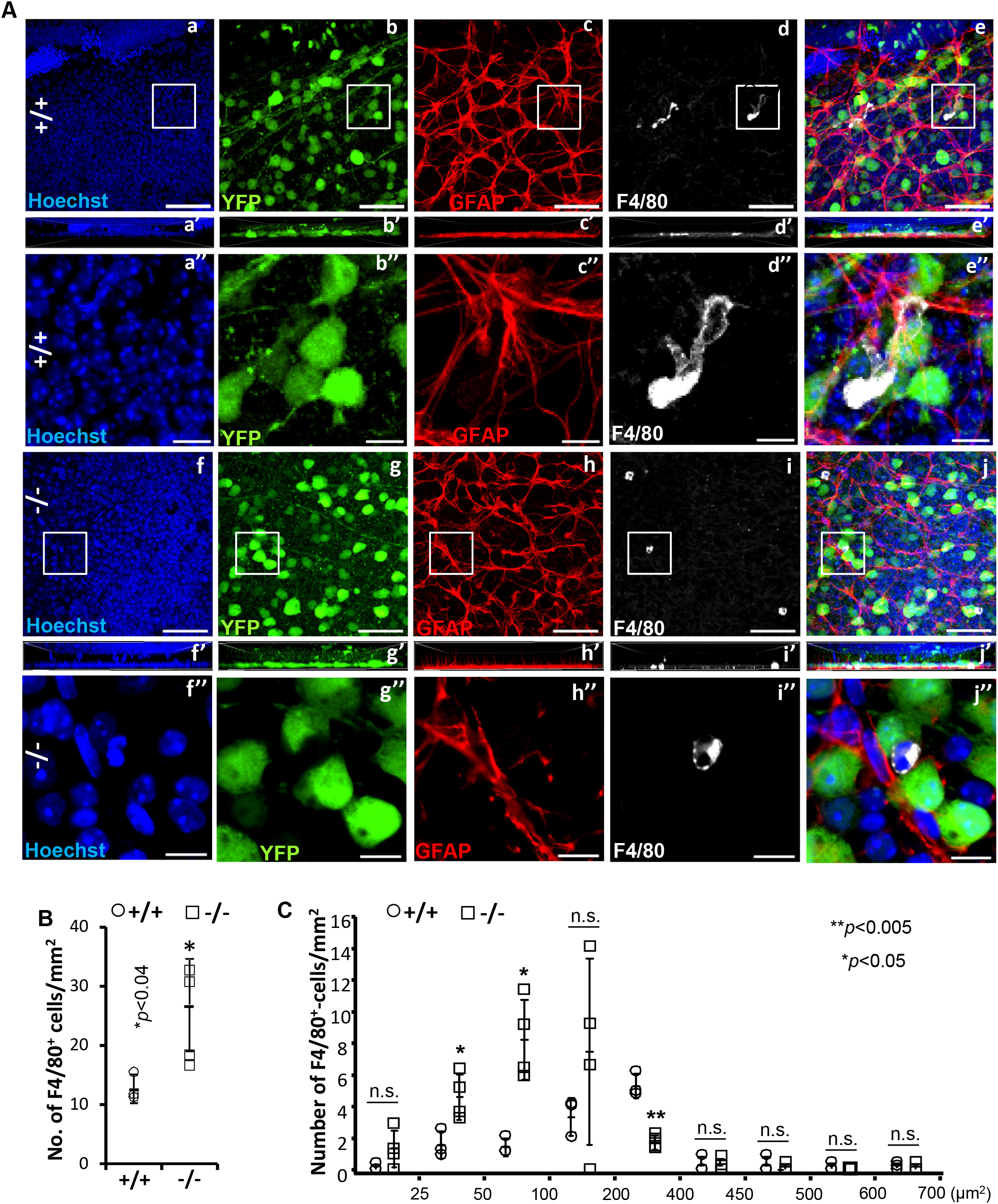
Activation of F4/80^+^-microglial in the ganglion cell layer of the retina by loss of *Ranbp2*. **A**. Confocal images of retinal flat mounts (ganglion neurons facing up) of +/+ and −/− mice immunostained for GFAP and F4/80 at d10. F4/80^+^-microglia in the retinae of +/+ mice present extended pseudopodial-like processes (a-e), whereas −/− have a significant increase of the number of activated and smaller F4/80^+^-microglia with an amoeboid morphology (f-j). There is no apparent gliosis between −/− and +/+ as reflected by the lack of changes in GFAP^+^-astrocytes between −/− (h’) and +/+ (c’). Images a’-j’ are lateral views of *z*-stacks of a-j and collapsed from a 25 μm thick-stack of 13 images captured 2μm apart. F4/80^+^-microglia of −/− show a clear and typical amoeboid morphology of activation (inset box in i is magnified in i’’), whereas resting F4/80^+^-microglia of +/+ show prominent morphological extended processes like lamellipodia/ pseudopodia (inset box in d is magnified in d’’). **B**. Quantitative analyses of F4/80^+^-microglia showing that there is a significant increase of the number of F4/80^+^-microglia in the ganglion cell layer of −/− compared to +/+ mice. Data are expressed as mean ± sd. Student’s *t*-test, *n*=3-4 mice/genotype. **C**. Quantitative analysis of the distribution of size (areas) of F4/80^+^-microglia between genotypes show that the ganglion cell layer of −/− mice have a significant shift of a pool of F4/80^+^-microglia with an average size of 400-450 μm² to a pool of F4/80^+^-microglia with an average size of 25-100 μm². Data are expressed as mean ± sd. Student’s *t*-test, *n*=3-4 mice/genotype. Scale bars=50 μm (a-j), 10μm (a’’-j’’); d10, day 10 post-tamoxifen administration; n.s., non-significant; −/−, *SLICK-H::Ranbp2^flox/flox^*; +/+, *SLICK-H::Ranbp2^+/+^*

### Loss of Ranbp2 causes the subcellular sequestration of Mmp-28 in YFP^+^-ganglion neurons of *SLICK-H::Ranbp2^flox/flox^*

Our studies have uncovered that Ranbp2 controls the proteostasis of Mmp11 and Mmp28 in a neural-type-dependent manner that culminate in non-cell autonomous responses [34,32,48]. For example, loss of Ranbp2 in Thy1^+^-motoneurons promotes the downregulation of Mmp28 in the sciatic nerve without affecting the secretion of Mmp28 from the soma of Thy1^+^-motoneurons [34]. Mmp28 is a regulator of macrophage activation and myelination [75–78]. In light of the foregoing findings, including that Mmp28 likely mediates neural-type selective paracrine responses upon loss of Ranbp2, we examined the effects of loss of Ranbp2 in YFP^+^-RGNs in the proteostasis and subcellular distribution of Mmp28. As shown in Fig 8A, there was a decrease of the levels of the 90 (dimer) and 48 kDa (monomer) isoforms of Mmp28 at d0 and d10, respectively, in the optic nerve, whereas the levels of the 90 kDa isoform of Mmp28 were transiently decreased in the retina at d0. Then, we examined the subcellular distribution of Mmp28 in YFP^+^-RGNs between genotypes. Mmp28 was found prominently in the interstitial space between YFP^+^-RGNs of wild-type mice, whereas there was widespread sequestration of Mmp28 in the intracellular compartment of YFP^+^-RGNs of *SLICK-H:: Ranbp2^flox/flox^* mice (Fig 8B).

**Figure 8.**
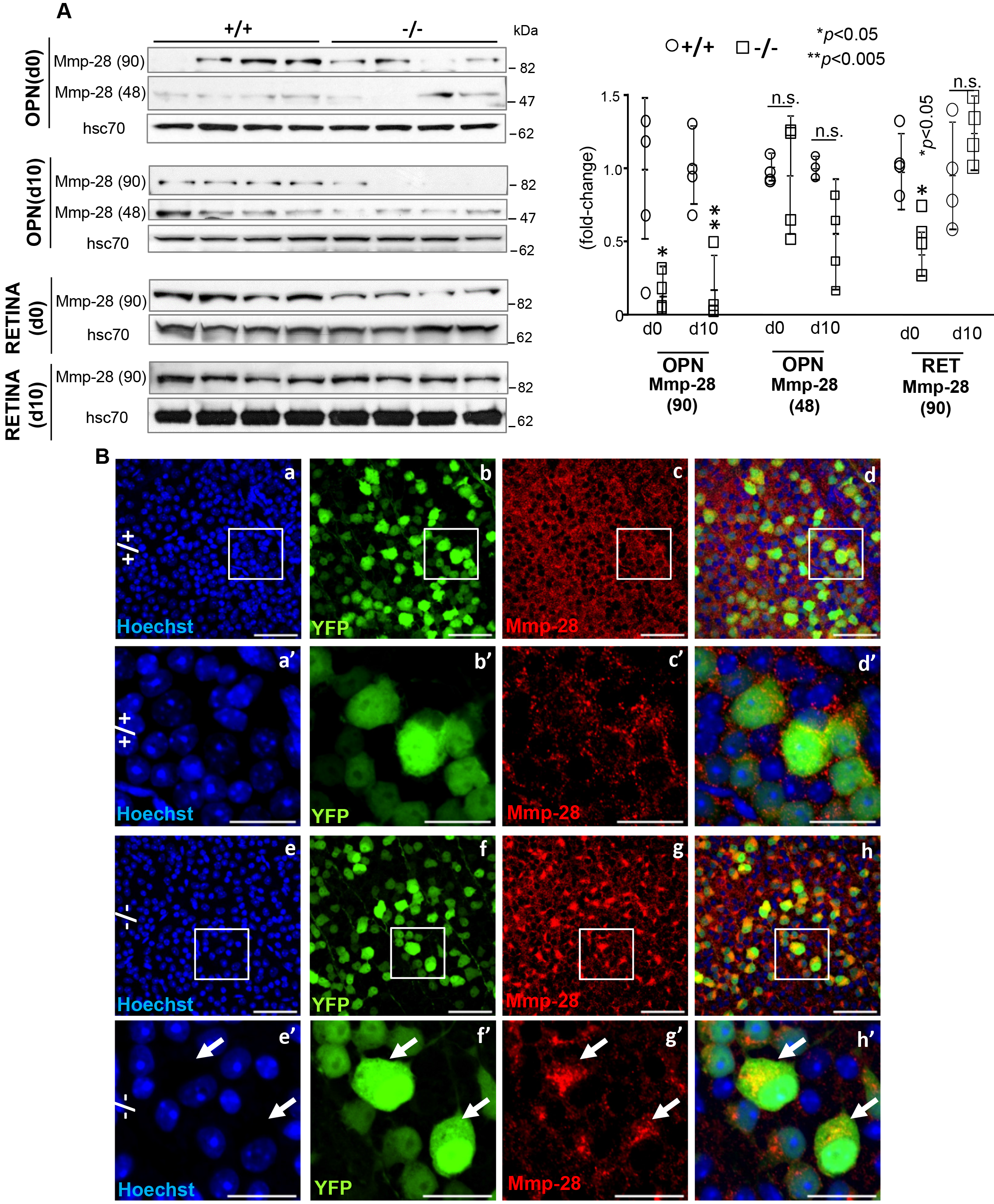
Suppression of proteostasis and biogenesis of Mmp-28 in the optic nerve and retinal ganglion neurons (RGNs) of the retina by loss of *Ranbp2*. **A.** Immunoblots (left) and quantitative analysis (right) of expression of the 90 (dimer) and 48 kDa isoforms of Mmp-28 in optic nerve (OPN) and retina (RET) at d0 and d10. In the optic nerve, −/− mice present an early decline in the 48kDa of Mmp-28 [Mmp-28(48)] at d0, whereas a decline in the 90kDa of Mmp-28 [Mmp-28(90)] is seen at d10. In retina, there is no expression of the Mmp-28(48) isoform and −/− mice have decreased levels of Mmp-28(90) at d0. Hsc70 is a loading control. Data are expressed as mean ± sd. Student’s *t*-test, *n*=4 mice/genotype. **B.** Confocal images of retinal flat mounts (ganglion neurons facing up) of +/+ (a-d, a’-d’) and −/− mice (e-h, e’-h’) immunostained for Mmp-28 at d10. Mmp-28 is localized outside the soma of YFP^+^-RGNs (extracellular space), whereas there is prominent sequestration of Mmp-28 in the perikarya of YFP^+^-RGNs of −/− mice (arrows). a’-d’ and e’-h’ are magnified images of inset regions of a-d and e-h, respectively. Scale bars= 50 (a-d, e-h) and 20 μm (a’-d’ and e’-h’). −/−, *SLICK-H::Ranbp2^flox/flox^*; +/+, *SLICK-H:: Ranbp2^+/++/+^*; hsc70, heat shock protein 70; d0 and d10 are days 0 and 10 post-tamoxifen administration, respectively; n.s., non-significant; OPN, optic nerve; RET, retina.

## Discussion

A growing body of evidence supports that dysregulation of nucleocytoplasmic trafficking is present in heterogeneous forms of sALS and fALS [1–9] and that the visual pathway, including RGNs, develop structural and functional impairments that may precede the development of ALS motor syndromes [6,42–45]. Ranbp2 is a unique vertebrate nucleoporin, which controls rate-limiting steps of nucleocytoplasmic transport. Ranbp2 is highly expressed in motoneurons and RGNs [34,35]. In the present study, a mouse model of ALS with rapid declines of motor performance that culminate in hind paralysis, respiratory distress and death, and without expression of Ranbp2 in Thy1^+^-motoneurons and RGNs, was used to examine the roles of Ranbp2 in Thy1^+^-RGNs. This disease mouse model represents a valuable tool to discern shared and unique molecular, cellular and pathobiological effects caused by loss of Ranbp2 between Thy1^+^-RGNs and those reported by a previous study in Thy1^+^-motoneurons [34].

The findings of this study support the following concepts. 1) RGNs and motoneurons share the dysregulation of nucleocytoplasmic transport by loss of *Ranbp*2. This disruption is manifested by the impairment of nucleocytoplasmic partition of Ran GTPase and nuclear import (e.g., importin-β) and export receptors (e.g., CRM1/exportin-1) and auxiliary substrates (e.g., HDAC4). 2) Ranbp2 is critical to RGN homeostasis; the somas of Thy1^+^-RGNs develop hypertrophy and their myelinated axons undergo a decline of axonal caliber following deletion of *Ranbp2*. These effects are accompanied by a functional delay of visual cortical responses to light stimuli. 3) With the exception of Ccl6, the optic nerve does not co-opt transcriptome changes with the sciatic nerve after loss of *Ranbp2* in RGNs and motoneurons. Loss of Ranbp2 in RGNs induces the pronounced and significant increase of the mRNA level of a critical immunomodulator, acetyl-CoA carboxylase 1 (Acc1), which is known to control *de novo* fatty acid synthesis and immunity in T-cells [69,70]. In naive mice lacking Ranbp2 in RGNs, the upregulation of Acc1 was accompanied by the multiphasic activation of microglia. 4) RGNs and motoneurons share the dysregulation of the proteostasis of Mmp28, which is another modulator of immunity and possibly myelination [75–78], but this effect likely arises by distinct mechanisms between RGNs and spinal motoneurons.

Like in motoneurons [34] and other cell types [32], the nucleocytoplasmic partitioning of the driver of nucleocytoplasmic transport, Ran GTPase, and nuclear import (importin-β) and export receptors (e.g., CRM1/exportin-1), were impaired in RGNs lacking Ranbp2 of *SLICK-H:: Ranbp2^flox/flox^*. Comparable disruptions were also observed in auxiliary substrates of Ranbp2 (e.g., HDAC4 and NR2F2). These observations are consistent with the regulation of rate-limiting steps of disassembly nuclear export cargoes and/or regeneration of importin-β for nuclear import by Ranbp2 [18,20–22,26]. Unexpectedly, the disruption of nucleocytoplasmic partition of these factors were accompanied by transcriptome changes in optic nerves that were distinct from those reported previously in the sciatic nerve [34]. These findings support a combinatorial coding mechanism whereby Ranbp2 regulates the nucleocytoplasmic transport of neural-type selective substrates and whose nuclear export or import is dependent on exportin-1, importin-β or both. It is unlikely that the nuclear import pathway plays a determinant pathophysiological role in motoneurons and RGNs lacking Ranbp2, because genetic complementation studies support that mice with loss of Ran-GTP and importin-β binding to RBD2 and RBD3 of Ranbp2 (e.g.; *Tg*-*Ranbp2^RBD2/-HA^* ::*SLICK-H::Ranbp2^flox/flox^*) rescue the motor deficits developed by mice with loss of *Ranbp2* in motoneurons [34]. This is in contrast with the severe degeneration of other cell types caused by losses of RBD2 and RBD3 functions of Ranbp2 (e.g., retinal pigment epithelium and developing cone photoreceptor neurons) [32]. A possible scenario is that impairment of a Ranbp2-mediated nuclear export pathway by docking of exportin-1 to the zinc-finger-rich domain of Ranbp2 [26] is impaired in RGNs (and motoneurons) lacking *Ranbp2* and that this impairment precludes the coupling of nuclear export with axonal transport of ribonucleic protein cargoes that are selective to RGNs or motoneurons. Motoneurons and RGNs are likely highly vulnerable to impairments in coupling of nuclear export with axonal transport because of the burden harbored by these neurons for the very long-distance transport of cargoes in axons that can reach a meter in size in some species (e.g. sciatic nerve). This concept is supported by several findings. First, haploinsufficiency or missense mutations in the nucleoporin, *Gle1*, which mediates nuclear export of mRNA, cause fetal motoneuron disease or ALS [79]. Second, the motor activities of the microtubule-based motor proteins, the kinesin-1 isoforms, KIF5B/KIF5C, which mediate the anterograde transport of selective cargoes, are regulated directly by Ranbp2 [80–83]. Third, Ranbp2 via its ZnF domain and docking of exportin-1 to ZnF potentiates the nuclear export of selective mRNAs and the translation of selective cargoes exported from the nucleus [26–28]. Finally, loss of Ranbp2 in motoneurons causes the pronounced increase of the levels of selective mRNAs and the decrease of their translation products that may result from the uncoupling of transport and translation of these mRNAs caused by loss of Ranbp2 [34]. Collectively, the data support the notion that the uncoupling of nuclear export of mRNA and protein cargoes from their axonal transport and/or translation by loss of Ranbp2 in RGNs may lead to the hypertrophy of their somata and axonopathy (e.g., axon atrophy).

Contemporary and recent studies indicate retinal involvement in ALS [6,42–45]. Specifically, these studies have shown that ALS is associated with structural changes by the thinning of the nerve fiber layer (e.g., unmyelinated axons of RGNs before they exit the optic cup), histopathological changes in inner retinal neurons, and functional deficits in sensory processing of the visual system measured by pattern reversal or luminance-evoked VEPs [6,42–45]. We have also found that mice with decreased levels of hnRNPA2B1 caused by selective loss of prolyl isomerase activity of Ranbp2 develop shorter latency of dark-adapted VEPs without changes in electroretinogram [33]. In this study, we found that impairment of nucleocytoplasmic transport by Ranbp2 was accompanied structurally by a decline of axonal diameter without loss of retinal ganglion cell bodies and functionally by a delay of the implicit times (latency) of VEPs before changes of motor behavior ensue [34], but without changes of VEP amplitudes. Hence, the normal VEP amplitudes support the absence of RGN loss, while the changes in latency of VEP indicate the functional impairment of transmission of the light stimulus to the primary or extrastriate occipitotemporal visual cortex [42]. This functional impairment resembles that seen in other retinal disease conditions, such as glaucoma, which is also linked to ALS, and in which there are delays in VEP timing.[84,85]

Recent studies have used transcriptome approaches to gain insights into the pathogenesis of ALS and the vulnerability of motoneurons to ALS [86]. In contrast to other ALS-causing RNA-binding proteins in which mutations purportedly promote broad transcriptome changes [87–89], knockdown of hnRNPA2B1 in the spinal cord led to the misregulation of a small number of transcripts (~20-30) [90]. Notably, little transcriptional overlap existed between the spinal cord and human iPSC-derived motor neurons [90]. Cultured spinal motoneurons expressing mutations in SOD1^G93A^ and TDP43^A315T^ showed few alterations in transcriptional expression and an overlap of only 20 transcripts that encoded proteins mainly with predicted extracellular localization [91]. Lumbar spinal motor and oculomotor neurons, which control eye movement and are spared in ALS, showed differences in broad transcriptional profiles of over 1,700 genes some of which presumably indicate a reduced vulnerability of oculomotor neurons to excitotoxicity by enhanced GABAergic transmission [92]. Finally, the transcriptome of microglia purified from spinal cord of ALS mice with SOD1^G93A^ compared to wild-type mice showed that ~2,000 transcripts are differential regulated by at least a 2-fold at late stages of the disease [86]. Collectively, these studies support the notion that distinct genetic insults in ALS promote heterogeneous transcriptional responses; however, the pathobiological relevance and the extent to which experimental and disease contexts contribute to apparent divergences of these transcriptional abnormalities remain unclear. By contrast, our study explored the effects of selective disruption of nucleocytoplasmic transport, which is shared by heterogeneous forms of sALS and fALS [1–6], in the transcriptome of the optic nerve. Further, we compared the transcriptome changes of the optic nerve with those previously reported in the same mice for the sciatic nerve [34].

Our analysis showed that apart from Ccl6 the transcriptomes of optic and sciatic nerves with concurrent loss of Ranbp2 in RGNs and motoneurons had no overlap in transcriptional changes. It is likely that the absence of such overlap reflects the distinct ontologies between oligodendrocytes (e.g., neural tube) and Schwann cells (e.g., neural crest) that ensheath with myelin the axons of optic and sciatic nerves, respectively [93]. In this respect, several transcripts, such as opalin (Tmem10) and Fam107a/Drr1, encode known glial markers of the CNS [94,95], while others, such as Ly6g6f, are up-regulated in frontotemporal lobar degeneration (FTLD) and linked to atrophy of the superior frontal gyrus, neuroinflammation and immunity [96,97] Notably, we found that the lipogenic enzyme, Acc1, was the target with the highest up-regulation in the optic nerve (~35-fold, *p* value < 0.05). Acc1 up-regulation is at the crux of a metabolic shift in the *de novo* fatty acid synthesis pathway by converting acetyl-CoA to malonyl-CoA, which is the carbon donor for the synthesis of long-chain fatty acids [67,68]. The stimulation of this pathway is critical to release the dependence of tumor cell growth and survival from exogenous lipids.[98] By contrast, Acc1 inhibition suppresses CD8^+^ T-cell expansion [69] and the genesis of T_H_17 cells with pathogenic roles in inflammatory and autoimmune diseases and favors instead the differentiation of anti-inflammatory T_reg_ cells, whose differentiation depends on exogenous fatty acids [70].

Our study found that in the CNS the pronounced up-regulation of Acc1 was linked to the multiphasic activation of microglia as manifested by the activation of ramified CD11b^+^ and CD45^+^-microglia, increase of F4/80^+^-microglia and development of amoeboid F4/80^+^-microglia intermingled between RGN of naive mice lacking Ranbp2 in RGNs. It is possible that an increase in dynamics between pseudopodial/lamellipodial and amoeboid F4/80^+^-microglia contributes to the increase of amoeboid F4/80^+^-microglia [74]. Current studies also indicate that these microglia represent resident microglia, which become activated by native insults instead of perivascular/systemic macrophages that typically show delayed recruitment triggered by nonnative insults that may compromise the brain-blood/retinal-blood barriers [74,99–102]. Regardless, our results suggest the activation of a paracrine signaling mechanism between RGNs and quiescent microglia resident in the retina that is dependent on the Acc1-dependent production of free-fatty acids by RGNs. This intercellular cross-talk is analogous to the chemokine-mediated paracrine and neuroglia signaling between axons of spinal motoneurons and surrounding Schwann cells triggered by loss of Ranbp2 in motoneurons [34]. In contrast to our findings in the sciatic nerve and other neurons however, we did not find changes in the levels of free fatty acids (octanoate and longer fatty acids) in the optic nerve [34]. The reason for this outcome needs further investigation. It is possible that the content of specific species of free fatty acids are affected by Acc1 in RGNs or oligodendrocytes. In this regard, metabolomics studies found that brains of haploinsufficient *Ranbp2* mice when challenged with the parkinsonian neurotoxin, MPTP, presented a decline of the medium-chain free fatty acids, pelargonate (9:0) and caprate (10:0), and the long-chain free fatty acid, arachidate (20:0) [53]. Alternatively, emerging studies indicate that suppression of Acc1 plays a fatty acid synthesis-independent role by promoting acetyl-CoA-dependent acetylation of proteins that trigger selective cellular responses. In this regard, it is noteworthy that both motor and RGNs share the nucleocytoplasmic deregulation of HDAC4, which is a substrate of Ranbp2 [34,33,64]. Hence, an increase of Acc1 levels combined with dysregulation of HDAC4 may result in a decline in the acetylation of selective substrates that ultimate contribute to microglia activation. Finally, since Ranbp2 and its ZnF domain potentiate the translation of selective transcripts exported from the nucleus [28,27], the up-regulation of *Acc1* may reflect a compensatory response or accumulation of *Acc1* transcripts arising from translation deficits of *Acc1* by loss of Ranbp2. This study will provide the foundation to elucidate between these mechanistic scenarios that are not mutually exclusive necessarily.

Recent data support the notion that selective metalloproteinases act directly as strong modifiers of disease progression in ALS and cytokine-mediated pathology [103,104]. Our prior studies have shown that Ranbp2 controls the proteostasis of selective metalloproteinases (e.g., Mmp11 and Mmp28) in a neural-type dependent manner and that these Mmps likely exert noncell autonomous effects in neighboring cells/tissues [32,34,48]. In motoneurons, Ranbp2 regulates Mmp28 proteostasis in the sciatic nerve but not in soma of motoneurons.[34] In this study and in accord to other studies, we found that Mmp28 is constitutively secreted to the interstitial space of the ganglion neural network of the retina. Mmp28 is a known regulator of responses of M1 and M2 type macrophages by mechanisms that remain elusive [76,75,105], but the physiological role(s) of Mmp28 in the CNS is unknown. Our studies indicate that Mmp28 is a strong candidate to regulate pro-signaling substrates in RGN-microglial signaling. Loss of Ranbp2 in RGNs caused the prominent and selective intracellular sequestration of Mmp28 in these neurons. The up-regulation of Acc1 may contribute to the Mmp28 sequestration by affecting the biogenesis of the endoplasmic reticulum and Golgi apparatus and membrane phospholipids [106,70]. However, ultrastructural analysis of soma of RGNs did not find overt ER/Golgi pathologies. It is possible that inhibition of the biogenesis and secretion of Mmp28 by loss of Ranbp2 in RGNs suppresses or promotes the release of anti-inflammatory or pro-inflammatory cytokines, respectively, from the interstitial space and that the homeostasis of selective cytokines contributes to microglial activation. Ccl6 is a chemokine whose production is strongly stimulated in inflammatory disorders [107,108] and its up-regulation is shared by motoneurons and RGNs upon loss of Ranbp2 [34]. Ccl6 acts as a macrophage chemoattractant [108] and is also expressed by microglia [109]. Since challenged macrophages lacking Mmp28 present enhanced chemotaxis and chemokine production [75,76], suppression of Mmp28 biogenesis in RGNs may promote Ccl6 stimulation to activate microglia and microglia-microglia signaling.

In summary, mechanistically our results support a model in which loss of Ranbp2 in RGNs promotes immune responses in the CNS that apart from Ccl6 and Mmp28 are distinct from the peripheral responses caused by loss of Ranbp2 in spinal motoneurons [34]. Our model indicates that Ranbp2 is a major regulator of *Acc1* expression and that they together play a central role in neuroimmunity and paracrine signaling between RGNs and microglia (Fig. 9). According to this model, up-regulation of *Acc1* by loss of Ranbp2 inhibits the biogenesis (e.g. secretion) of Mmp28 and stimulates the release of selective chemokines (e.g. Ccl6) and/or fatty acids that promote neuroglial signaling and microglial priming and activation. These effects are accompanied by the biphasic regulation of the cell cycle protein, Pttg1/securin, and its potential transcriptional target, Cdnk1 (p21)[66], and of stress-signaling, cell migration and cell surface membrane markers, such as, Ly6g6f and Fam107a [95,97,110–112]. Hence, these targets are strong candidates to modulate the proliferation and migration of microglia. Although microglial dysfunction is linked to ALS [113], emerging and contradictory data also suggest that there is a lack of convergence in neuroinflammation between fALS and sALS [114]. Our data indicate that disruption of nucleocytoplasmic transport shared by fALS and sALS, and between retinal ganglion and motor neurons, promotes unique endophenotypes in RGNs and microglia and that the visual pathway may offer pathognomonic signs of ALS and/or other motoneuron diseases. These molecular and cellular mechanisms deserve heightened attention because nucleocytoplasmic transport is also disrupted in other neurodegenerative diseases [10–13].

**Figure 9.**
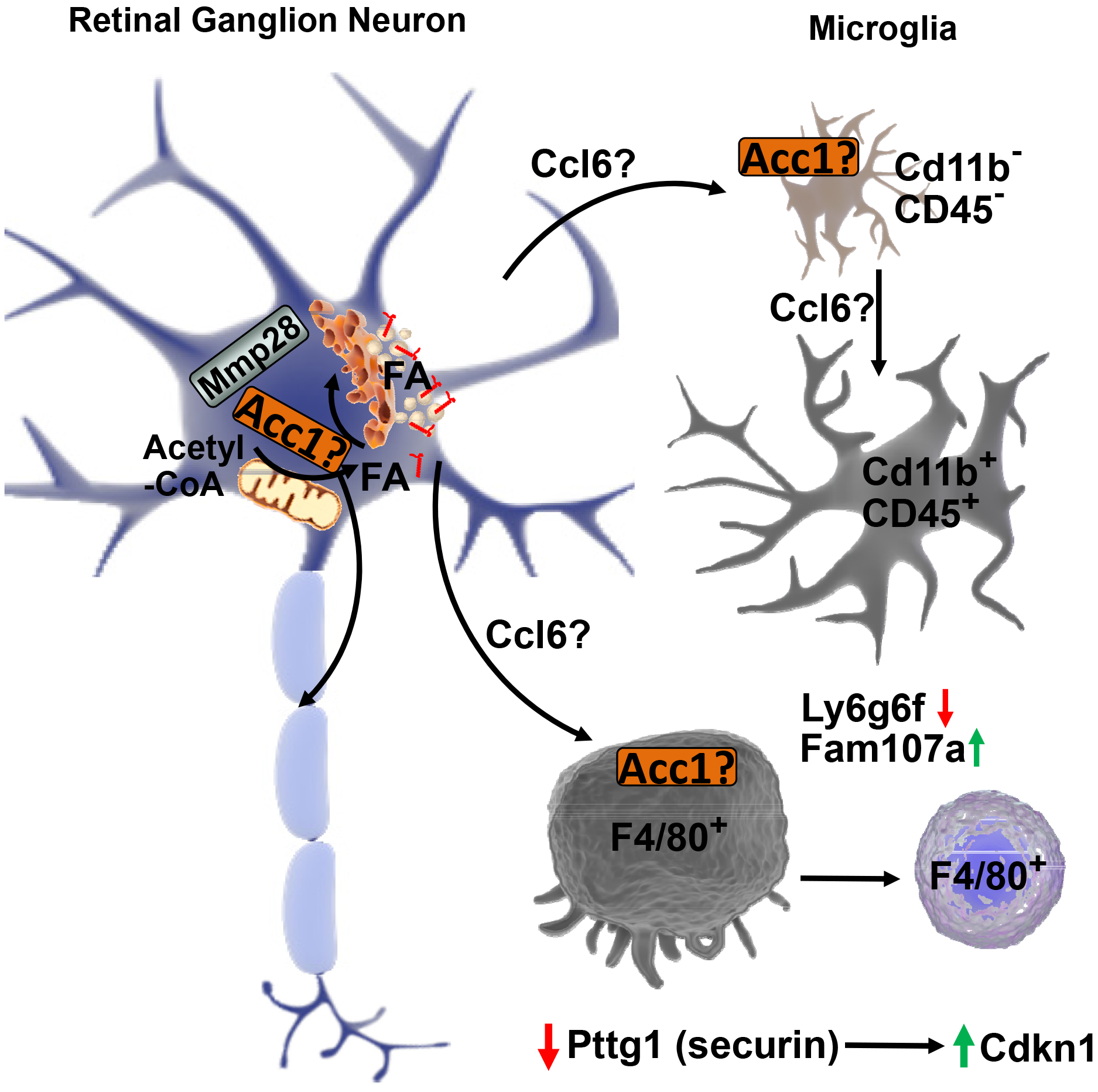
Mechanistic model of microglial activation by loss of Ranbp2 in Thy1-retinal ganglion neurons (RGNs). Loss of *Ranbp2* in RGNs triggers the up-regulation of *Acc1* and the stimulation of free fatty acids and/or Ccl6 in RGNs, microglia or both. There is also intracellular sequestration of Mmp28 in RGN that relieves the suppression of Ccl6 production and stimulates neuroglia signaling and the multiphasic activation of microglia with Cd11b^+^ and CD45^+^-ramified and F4/80^+^-amoeboid morphologies. These effects are also accompanied by changes in expression of cell-cycle and surface antigen/cell migration regulators, such as the downregulation of Pttg1 and Ly6g6f and the up-regulation of Cdkn1 and Fam107a in RGNs and/or microglia (see text for details).

## Author contributions

KC and PF conceived and supervised the study; KC, NP and PF designed experiments; KC, MY and DY performed experiments; PF provided new tools and reagents; KC, DY, NP and PF analyzed data; KC and PF wrote the manuscript.

## Acknowledgments

We thank Guoping Feng (MIT, Cambridge, MA) for SLICK-H mice, Ian Macara (Vanderbilt University, Nashville, TN) for the antibody against Ran-GTP, Sandra Stinnett for help with statistical analyses of axonal morphometry (Duke University, Durham, NC), Ying Hao for help with the processing of the specimens for transmission electron microscopy (Duke University, Durham, NC).

## Availability of data and materials

The datasets supporting the conclusions of this article are available in the Sequence Reads Archives (SRA; https://www.ncbi.nlm.nih.gov/sra) of the National Center for Biotechnology Information (NCBI) with the accession number: SRP139153. Datasets of a total of ∼17.4 billion bases of unprocessed RNA sequencing were deposited as FASTq files at the SRA. The project overview was deposited with the Bioproject accession number: PRJNA449172. The optic nerves of *SLICK-H::Ranbp2^+/+^*, *SLICK-H::Ranbp2^flox/flox^* and *Tg-RBD^2/3*-HA^::SLICK-H::Ranbp2^flox/flox^* have the respective biosamples accession numbers: SAMN08891694, SAMN08891693 and SAMN08891695. Other datasets and materials used and/or analyzed during the current study are available from the corresponding author on reasonable request.

---

**Supplemental Figure 1.**
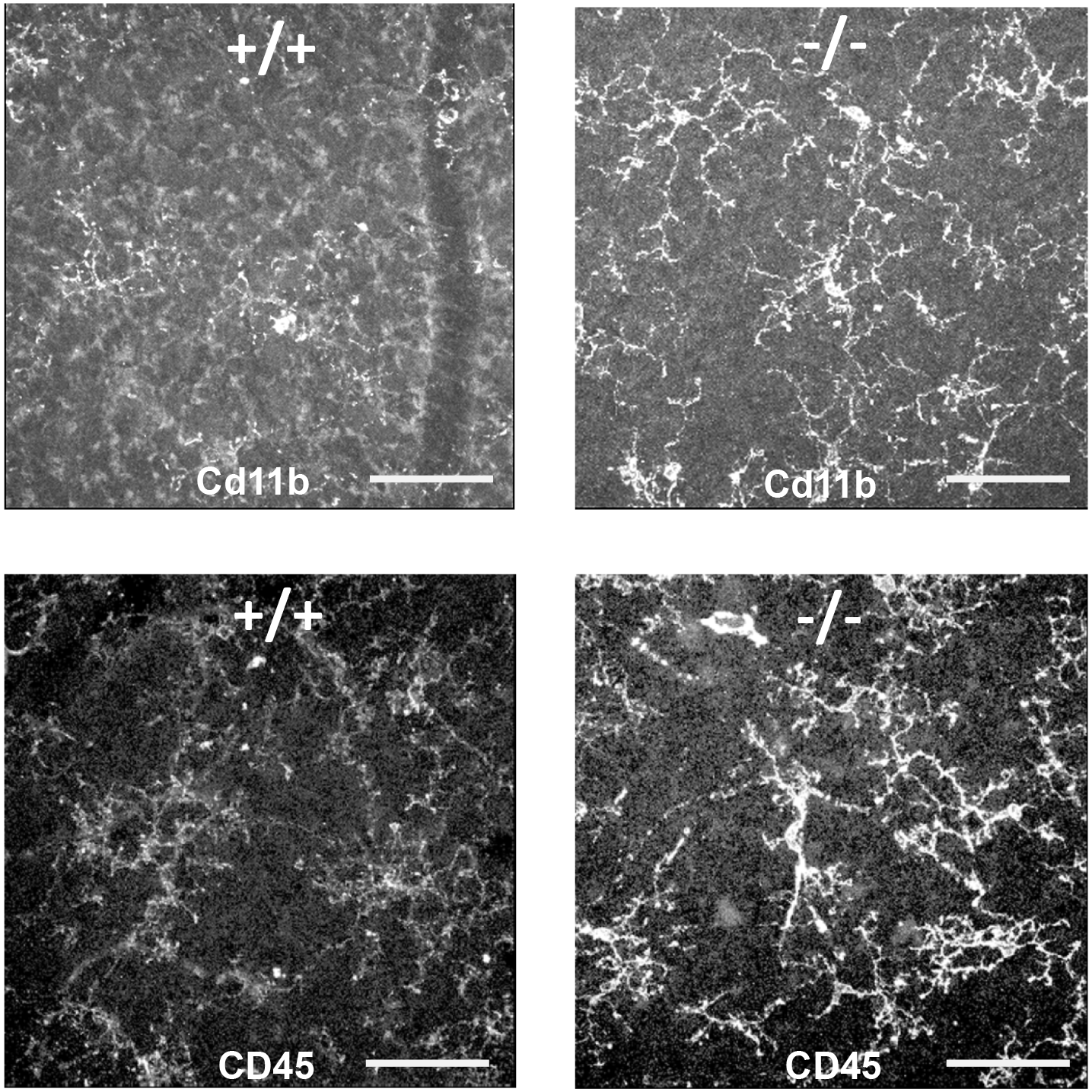
Activation of ramified CD11b^+^ and CD45^+^-microglia in the retinal ganglion cell layer caused by loss of *Ranbp2* in retinal ganglion neurons (RGNs). Confocal images of retinal flat mounts (ganglion neurons facing up) of +/+ and −/− mice immunostained for CD11b (**A**) and Cd45 (**B**) at d10 showing the activation of ramified microglia in −/− compared to +/+ mice. In comparison to +/+, −/− mice show stronger immunolabeled CD11b^+^ and CD45^+^-microglia. Image panels of Fig 6 underwent identical post-acquisition gamma-intensity corrections for the sole purpose of showing that weakly immunostained microglia of +/+ mice have also ramified morphology. Scale bars=50μm; d10, day 10 post-tamoxifen administration; −/−, *SLICK-H::Ranbp2^flox/flox^; +/+*, SLICK-H::Ranbp2^+/+^.

